# QTL mapping reveals genomic regions for yield based on incremental tolerance index to drought stress and related agronomic traits in canola

**DOI:** 10.1101/2020.01.06.896688

**Authors:** Harsh Raman, Rosy Raman, Ky Mathews, Simon Diffey, Phil Salisbury

## Abstract

Drought stress especially at the reproductive stage is a major limiting factor that compromises the productivity and profitability of canola in many regions of the world. Improved genetics for drought tolerance would enable the identification and development of resilient varieties, resulting in increased canola production. The main objective of this study was to dissect the genetic basis of seed yield under water-limited conditions in canola. A doubled haploid population derived from a cross between two Australian parental lines, RP04 and Ag-Outback, was evaluated to identify the genetic variation in fractional normalised deviation vegetative index (NDVI), above ground shoot biomass accumulation, flowering time, and plasticity in seed yield under irrigated and rainfed field conditions in two consecutive years. An irrigation treatment was applied at the 50% flowering stage and an incremental drought tolerance index (DTI) was estimated for seed yield. By utilising a genetic linkage map based on 18,851 genome-wide DArTseq markers, we identified 25 genomic regions significantly associated with different traits (LOD ≥ 3), accounting for 5.5 to 22.3% of the genotypic variance. Three significant genomic regions on chromosome A06, A10 and C04 were associated with DTI for seed yield. Some of the QTL were localised in the close proximity of candidates genes involved in traits contributing to drought escape and drought avoidance mechanisms, including *FLOWERING LOCUS T* (*FT*) and *FLOWERING LOCUS C* (*FLC*). Trait-marker associations identified herein can be validated across diverse environments, and the sequence based markers may be used in a marker assisted selection breeding strategy to enhance drought tolerance in canola breeding germplasm.

## Introduction

Oilseed rape (*Brassica napus* L) is one of the major crops grown for vegetable oil production and is grown on 57 million hectares worldwide. Climate change accompanied by the prevalence of elevated temperatures and variation in weather patterns poses a great challenge to canola production and profitability. It is estimated that 30% more oil crop production is required by 2050 to meet growing demands for vegetable oil, biodiesel and stock feed markets [1]. This projected increase in productivity has to come largely from improved genetics, advanced genetic selection technologies and improved agronomic practices on farm. In Australia, canola is grown over ∼2.5 million ha, however the area sown depends upon seasonal rainfall and global price index (www.australianoilseeds.com/). It is sown in autumn (March-May) and harvested in early summer months (November-Dec). During the 7 month crop season, provision of an adequate water supply (either by rainfall or irrigation) is critical for canola growth, especially at the flowering and pod filling stages. Drought stress at these stages (mid and terminal drought) not only adversely affects seed productivity, but also reduces the oil content and alters fatty acid composition [2–4]. Development of canola varieties having more plasticity to cope with water shortages would be useful for improving yield in drought-prone environments.

Natural genetic variation for drought tolerance exists in several native grass species, and some agricultural crops [5]. These plant species cope with drought stress via various mechanisms such as ‘escape’ - accelerating reproduction processes such as early flowering and seed set [6] and or via ‘tolerance’ mechanisms involving water use efficiency [7–10]. Early establishment (vigour) in canola ensures sufficient plant biomass is available to produce reproductive structures such as flowers and pods. It is particularly important if higher yields are to be achieved in drought stress conditions. Highly vigorous plants are likely to provide better ground-cover leading to greater stored soil moisture, which is reported to increase water-use efficiency [11] as well as weed control [12]. Early plant vigour has been measured using biomass cuts, leaf area index and digital imaging [13–15]. It has also been shown that normalised difference vegetation index (NDVI) can be used as a surrogate for biomass accumulation, fractional ground cover and plant vigour in canola [16–18].

Canola breeding programs have been developing varieties with a range of traits such as early vigour, flowering time and resistance to diseases. While the genetic basis of flowering time and resistance to various diseases is reasonably well understood in canola; limited knowledge is available on the component traits that contribute to improved yield under drought stress (drought tolerance). In addition, biotic and other abiotic stresses e.g. heat and soil acidity interact with drought stress, result in a complex system where it is challenging to dissect the genetic basis of drought tolerance. As a consequence, it has been difficult to make significant genetic progress in improving seed yield under drought stress conditions through conventional breeding methods. Previous studies have utilised the ‘yield sensitivity’, the difference between yield between irrigated and DRY environments in *B. napus* [19] and *B. rapa* [20], as a measure of drought tolerance. Genetic loci associated with drought tolerance have not been identified in Australian canola germplasm so far.

In this study, we implement a statistical model based on an incremental crop tolerance approach [12] to investigate the genetic basis of yield in response to drought stress at the reproductive stage (terminal drought). This approach relies on an appropriate experimental design with drought and irrigated treatments properly replicated. A drought tolerance index (DTI) is then calculated as the deviation from the regression of the drought treatment against the irrigated treatment. Following a quantitative trait locus (QTL) analysis approach, we identified several genomic regions for productivity traits; fractional ground cover, shoot biomass accumulation, flowering time and seed yield; measured as DTI in canola. Our results showed that DTI in terms of seed yield is a complex trait controlled by several QTL and plant architectural QTL may have pleiotropic effects on canola yield.

## Material and methods

### Plant materials

A doubled haploid population (n = 165) derived from an F_1_ hybrid developed from two Australian parental lines; RP04 and Ag-Outback, designated as ‘RADH’ was constructed at the Agriculture Victoria, Horsham, Australia and used in this study. This population segregates for both qualitative and quantitative resistance to the fungal pathogen, *Leptosphaeria maculans*, causing blackleg disease [21]. Both parental lines, RP04 and Ag-Outback are also shown to differ in fractional ground cover [17]. However, genetic loci associated with fractional ground cover have not been identified.

### Experimental field design

A total of 188 lines comprising 165 RADH lines along with both parental lines (RP04 and Ag-Outback) and 21 check cultivars; 44C73, 46C76, Ag−Castle−4, Ag-Outback, Av-Opal, Av-Sapphire, BLN3303−CR0302, BLN2762, BLN3614, DHC2211, DHC2261, Drakkar, RP012*S, RR005, SturtTT, Surpass400, Topas, Zhongshu−Ang−No4, Zhongyou−821, ZY001 and ZY011 were grown as a split-plot design with two treatments; control (irrigated) and drought (rainfed) conditions during the optimum cropping season (April to December). Two experiments were carried out; experiment-I was sown on 2^nd^ May 2014 at the Wagga Wagga Agricultural Institute (WWAI, Wagga Wagga, NSW, Australia; latitude: −34.930°, longitude: 147.767°) while, experiment-II was sown on 7^th^ May 2015 at WWAI. The split-plot experimental design structure was the same for both experiments with a total layout of 48 Rows by 16 Ranges. The 2 replicates (Rep), were aligned along the Rows (1-24 and 25-48), as were the 2 Main Plots (Rows 1-12 and 13-24 within Rep I, and Rows 25-36 and 37-48 in Rep II). The treatment (Trt) with two levels (irrigated and rainfed) were randomised to each Main Plot in a restricted manner such that the irrigated treatment was allocated to the two inside Main Plots (Rows 13-24 (Rep I) and 25-36 (Rep II) for logistical reasons, and is the same for both years (Supplementary Figure 1.1). All entries were randomly allocated to a Sub-plot (intersection of a Row and Range) within a Main Plot, except SturtTT which was randomly allocated to six Sub-plots within each Main Plot. The layout for the controls is shown to illustrate the randomisation which is different between years (Supplementary Figure 1.2). For the irrigated treatment, supplementary irrigations (two in 2014 and one in 2015) were provided at the flowering stage (50%) in order to maximise the difference between treatments.

For both experiments 1,400 seeds of parental lines and their DH lines were sown in plots (2 m wide x 10 m long) with an eight-row cone-seeder. Plots were cut-back to 8 m at BBCH growth stage 40 by foliar application of Glyphosate (round-up) herbicide with a shielded spray boom. All plots were sown with granular fertiliser (N: P: K: S, 22:1:0:15) at the rate of 150 kg ha^-1^. The fertiliser was treated with Jubilee fungicide (a. i. flutriafol at the recommended rate) to protect from *L. maculans*, a fungus causing blackleg disease. At the 4-6 leaf stage, the plants were sprayed with Prosaro ® 420C (Bayer) to protect from blackleg and the second application was done at the flowering stage to control sclerotinia stem rot. Temperature and rainfall are summarised for each experiment in Supplementary Figure 1.

### Trait measurements

Four traits measured in both years were: fractional ground cover, shoot biomass accumulation, flowering time and seed yield. Fractional ground cover was assessed using the normalised difference vegetation index (NDVI) from 40 days of sowing at 7-10 days interval until flower initiation. This was measured from whole plots using a hand held device, the GreenSeeker® (NTech Industries Inc., Ukiah, CA, USA) following manufacturer’s instructions. For above ground shoot biomass, five (2014) to ten (2015) plants of parental lines and their DH lines, and check varieties from all replicated plots were cut from the hypocotyl/root junction with steak knives at BBCH growth stage 40 stage (weeks after sowing). Samples were taken from the middle two rows early in the morning to avoid drying-out, and transported to a laboratory, where samples were weighed on a two decimal scale balance. Shoot biomass was expressed in g/plant on fresh weight basis. Days to flowering were recorded at two stages: first flowering-when 25% of the plants had at least one flower open and last flowering; when all plants (>95%) show at least one flower. Flowering time was assessed three times per week. Plots were harvested with a plot header (Kingaroy, Australia) at full maturity (1st wk of December), seed were cleaned with Kimseed and then weighed in a laboratory. Yield was estimated and expressed in t ha-1.

### Genotypic data and physical mapping of marker associations

A genetic linkage map based on 2,317 binned markers (representing to 6,442 DArTseq SNPs and 12,409 in silico DArTs; Raman et al 2019), covering a genome-wide length of 3,118cM was used to identify trait-marker associations. Alignment of the genetically mapped markers to the reference *B. napus* physical genome of cv. Darmor-bzh [22] was performed using BLASTn analysis (e-value <1e-20). The physical location of the candidate genes was downloaded from annotated *B. napus* gene models genome assembly (http://www.genoscope.cns.fr/blat-server/cgi-bin/colza/webBlat) and further used to determine the physical distance between the candidate *B. napus* genes and significant markers.

### Statistical and QTL analyses

The drought tolerance index (DTI) was developed using a regression approach [12]. The regression approach defines the DTI as the deviation from the regression of the genotype effects for a given trait under rainfed conditions on genotype effects under irrigated conditions, that is, *E* in the following linear regression equation,

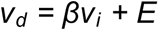

where *v_d_* is the vector of genotype effects for seed yield with drought treatment; *v_i_* is the vector of genotype effects for seed yield with the irrigated treatment; *β* is the slope of the regression, estimated by the s_di_/s_i_, where s_di_ is the genetic covariance between drought and irrigated and s_i_ is the genetic variance of the irrigated treatment. This regression model was incorporated into the linear mixed model for the treatment by genotype effects using a reduced rank model of order 1 plus a scaled identity (diagonal) matrix for the genotype effects for the irrigated treatment.

The QTL analysis for all traits proceeded as described in Raman et al. [23] and combines approaches presented previously [24]. Only yield was evaluated using the DTI approach as this was measured after the irrigated treatment was applied at the first flowering stage. There are three steps in the model fitting process: determination of a ‘baseline’ model, a genome scan to test each marker and the marker selection stage where the final model is determined. In the first step, a phenotypic (non-marker) analysis is performed for the DTI of each trait and any global, local or extraneous spatial variation are modelled following the approach of Gilmour et al. [25] and using the diagnostics presented in Stefanova et al. [26]. Broad-sense heritability and treatment by genotype best linear unbiased predictors are provided from this model. In the next step, each marker-trait association is tested via a genome scan where each marker, i = 1, 2, …, 2317 is fitted to the baseline model as fixed effects and the remaining markers incorporated into the model as a random effect. In the final marker selection steps, markers with a P-value <0.05 from a Wald test of the fixed marker effect from the genome scan step are retained as candidate markers. Co-linear (co-segregating) markers and/or markers in close proximity (<20cM) are discarded. Remaining candidate markers are fitted as fixed effects (whilst all other markers are fitted as random effects) and backward selection is performed to identify statistically significant markers (declared as putative QTL) and arrive at a final model. The LOD scores, percentage genotypic variation explained and the additive effects of significant markers are computed fusing the final model. All analyses were conducted in R [27] using the ASReml v3.0 library [28]. For simplicity, QTL identified in the genomic regions which were mapped within 0.5cM (∼550 kb) of the *B. napus* genome (∼1Gb) were designated as the same QTL.

## RESULTS

### Genetic variation in vegetative and reproductive traits

Genotype by treatment interaction for different vegetative traits and seed yield was initially investigated in the RADH population. A likelihood ratio test comparing models with (full model) and without (reduced model) the genotype by treatment interaction showed that this interaction was significant for full flowering (100%) and seed yield in both consecutive years of phenotyping. Mean yield clearly showed that both environments (2014 and 2015) were different (Table 1, Figure 1). This was supported by the meteorological data; total rainfall varied between years, with 2015 being a drought year (620 mm) and above average rainfall during the 2014 season (460mm) Supplementary Figure 1. As expected, values for seed yield were higher under the WET (irrigated) treatment compared to DRY (rainfed), including in check genotypes (Figure 2). For example, the mean seed yield for WET in 2015 was 2.2ha-1 compared to 1.7tha-1 for DRY, with a similar but less distinct difference in 2014 (Figure 1). A significant interaction indicates a re-ranking of genotypes between the WET and DRY treatments. Among the parental lines of the RADH population and the check genotypes, the maximum yield reduction occurred in Topas (vernalisation responsive) while the minimum yield reduction was in AV-Opal, followed by Drakkar. These results showed that the maternal parent, RP04 is more sensitive to drought stress than Ag-Outback (Figure 2).

**Figure 1:**
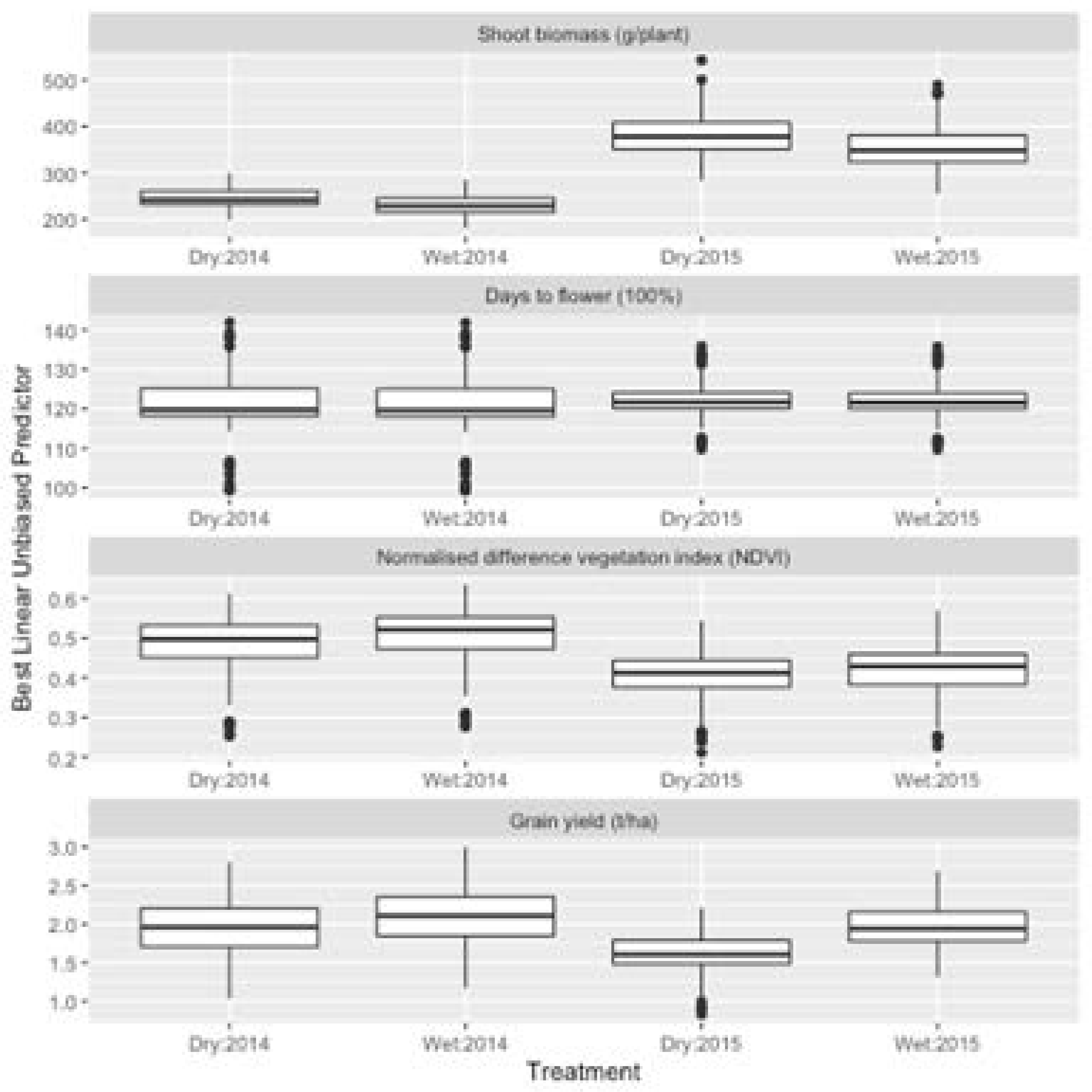
Box plots showing genetic variation for shoot biomass (g/plant), days to flower (100%), normalised deviation vegetative index (NDVI) and grain yield (t/ha) in the doubled haploid population derived from RP04/Ag-Outback evaluated under field conditions Best Linear Unbiased Predictors (BLUPs) were plotted for each treatment (WET and DRY) across 2014 and 2015 experiments. DH population means are indicated as solid lines. Vertical bars represent the range of trait variation within the DH population.

**Figure 2:**
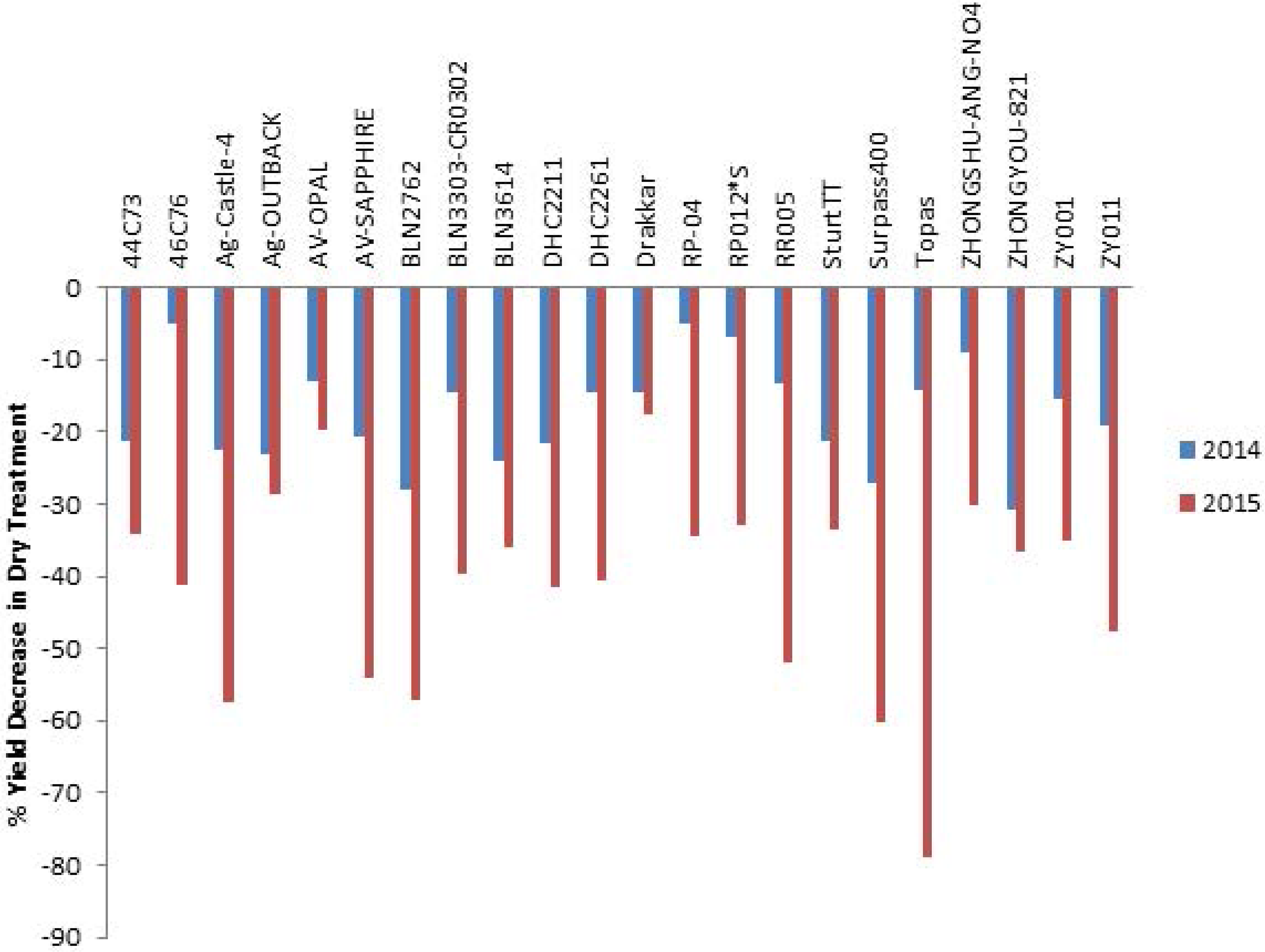
Per cent yield decrease in check and parental lines of RADH population derived from RP04/Ag-Outback.

**Table 1:** Phenotypic means for traits evaluated in a DH population from RP04/Ag-Outback

| Year | Trait | Minimum | Maximum | Mean | *CV | Reliability (h<sup>2</sup>, %) | Accuracy (%) |
| --- | --- | --- | --- | --- | --- | --- | --- |
| 2014 | First flowering | 87.59 | 131.08 | 107 | 2.7 | 97 | 98.5 |
| 2015 | First flowering | 231.13 | 255.64 | 236.7 | 0.7 | 96.6 | 98.3 |
| 2014 | Last flowering | 102.36 | 141.61 | 121 | 2.3 | 95.6 | 97.8 |
| 2015 | Last flowering | 241.76 | 263.73 | 249.2 | 0.8 | 93.7 | 96.8 |
| 2014 | NDVI (27th June) | 0.29 | 0.63 | 0.5 | 8.1 | 92.3 | 96.1 |
| 2015 | NDVI (11th August) | 0.48 | 0.77 | 0.6 | 7.4 | 69.1 | 83.1 |
| 2015 | NDVI (14th July) | 0.18 | 0.41 | 0.3 | 14.9 | 74.4 | 86.3 |
| 2014 | NDVI (20th June) | 0.23 | 0.52 | 0.4 | 9.7 | 91.7 | 95.8 |
| 2015 | NDVI (27th July) | 0.25 | 0.59 | 0.4 | 13.2 | 71.1 | 84.3 |
| 2014 | NDVI (8th July) | 0.45 | 0.71 | 0.6 | 8.8 | 73.2 | 85.6 |
| 2014 | Shoot Biomass | 217.19 | 299.22 | 238 | 25.2 | 36.7 | 60.6 |
| 2015 | Shoot Biomass | 327.03 | 504.73 | 367 | 25.1 | 29.6 | 54.4 |
| 2014 | Yield (t ha<sup>-1</sup>) | 0.67 | 3.53 | 2.03 | - | 93.2 | - |
| 2015 | Yield (t ha<sup>-1</sup>) | 0.44 | 2.96 | 1.8 | - | 71.3 | - |

\*CV: Coefficient of variance

The ranges of trait values for the DH lines were generally beyond the values of the two parents, suggesting transgressive segregation (Supplementary Figure 2). Broad sense heritability (*h*^2^) of different traits was used to quantify the percentage of total phenotypic variation explained by the genotypic component (Table 1). Moderate to high *h^2^* values were observed for flowering time (*h^2^* = 93.7 to 97%) in both environments, whereas shoot biomass, measured at early vegetative stage, had the lowest *h^2^* (29.6 to 36.7%; Table 1), indicating that shoot biomass measurements were not as reliable. High *h*^2^ values indicate that the genetic analyses of traits investigated in this study were reliable and suitable for genetic analyses.

We are also interested in the relationship between different traits both across and within years and describe these using pair-wise correlations (Figure 3, Supplementary Table 1). As expected, there were high genetic correlations between NDVI measurements and plant establishment across years which varied from 0.75 to 0.76 (2014) and 0.69 to 0.74 in 2015 (Supplementary Table 1). Within both years, high genetic correlations between NDVI measurements taken periodically (*r* = 0.9 to 1 in 2014, and *r* = 0.85 to 1 in 2015, Supplementary Table 1) were observed, suggesting that NDVI measured after 30 days of plant emergence (6 wk after sowing) provides a good estimate for predicting early vigour in canola. We also observed high genetic correlations for first (25%) and last flowering (100%) (*r* = 0.92 to 1.00 in 2014 and *r* = 0.96 to 1.00 in 2015, Supplementary Table 1). Trait correlation varied across both years (Figure 3); there was high correlation for NDVI (0.75), followed by seed yield (*r* = 0.73); moderate correlation for shoot biomass (*r* = 0.38) and poor correlation for flowering (*r* = 0.15). Genotypic variation in NDVI showed moderate correlation with seed yield (*r* = 0.43) across both 2014 and 2015 environments, while biomass accumulation had a poor correlation with seed yield (*r* = 0.11 to 0.14). Plant architectural traits; plant height, and early vigour showed moderate positive correlations with seed yield (*r* = 0.4 to 0.5) across both environments (Supplementary Table 1).

**Figure 3:**
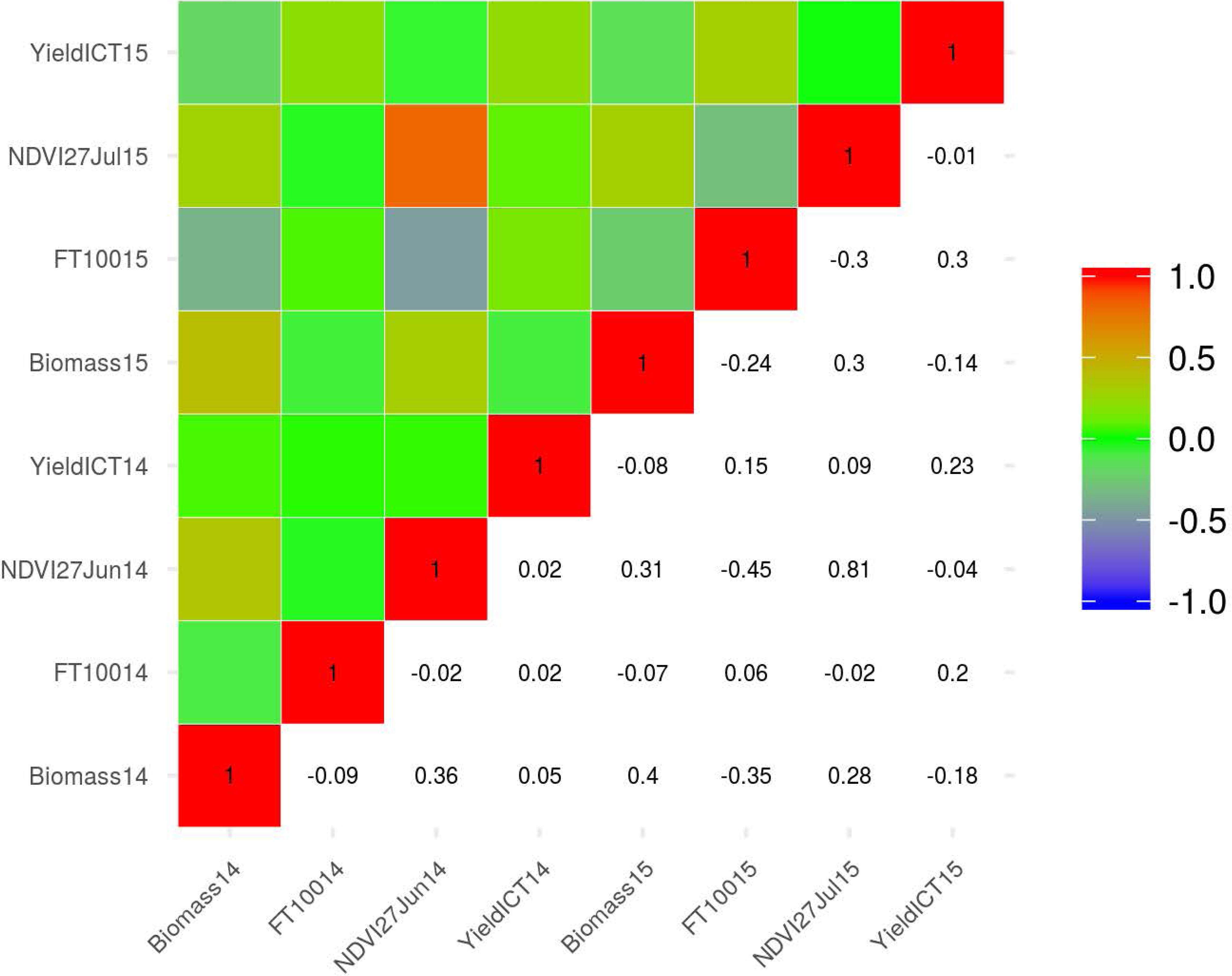
Pair-plots showing Pearson’s correlation coefficient (r) for shoot biomass (g/plant), days to flower (100%), NDVI and grain yield (t/ha) evaluated in field plots (20 sq M) of DH lines. Details are given in Supplementary Table 1.

### Identification of DH lines with significant additive allelic effect

To determine genetic variation for response to drought stress among DH lines, we plotted the predicted (marker) additive genetic effects for DH lines for WET treatment against those for DRY treatment. Several lines showed significant variation in the DTI (Figure 4). For example, DH lines 05CCD-0312, 05CCD-0342 and 05CCD-0369 had above average additive effects for drought tolerance, the latter in both years; whilst 05CCD-0225, 05CCD-0370, 05CCD-0286 and 05CCD-0203 DH lines had the below average additive effects.

**Figure 4:**
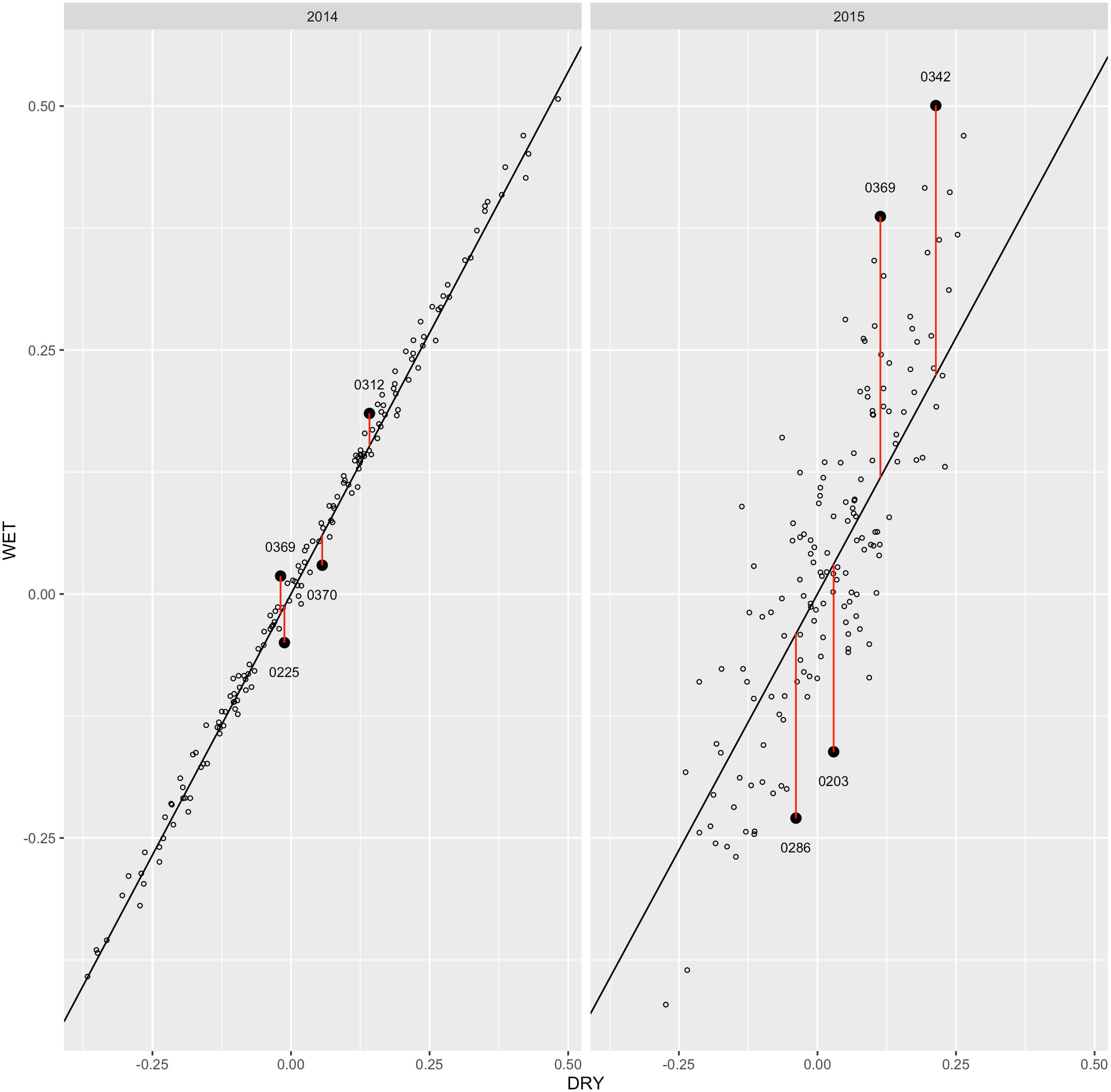
Predicted marker additive genetic effects for DH lines for WET treatment plotted against those for DRY treatment for each year. The genetic regression line is shown for each year and DH lines with large deviations from regression are identified with a solid circle and labelled with their DH index (last 4 digits of their full name, i.e. 05CCD has been dropped).

### QTL for yield measured using drought tolerance index

By using the DTI approach, 9 QTL (3 significant and 6 suggestive QTL) were identified for seed yield on chromosomes A03, A06, A09, A10, C03, C04, C08 and C09 (Table 2). These QTL accounted for 1.9% (C09) to 7.4% (A10) of the genotypic variance (R^2^). All QTL identified herein could explain a total of 36.8% of the R^2^ collectively across environments. Both parental alleles contributed to increase DTI for seed yield in the DH population from RP004/Ag-Outback (Table 2) However approximately 50% of the allelic effects for significant QTL were from the paternal parent of RADH population, Ag-Outback.

**Table 2.** QTL associated with drought tolerance index for seed yield in a DH population from RP04/Ag-Outback. DTI was calculated as described previously (Lemerle et al. 2006).

| Phenotyping year | Marker name | Chromosome | Physical position (bp) | Allelic effect (t/ha) | Probability (P =0.05) | LOD score | Genotypic variance explained (%) |
| --- | --- | --- | --- | --- | --- | --- | --- |
| *Statistical significant QTL* |  |  |  |  |  |  |  |
| 2014 | 31312620:A>G | A06 | 19833237 | -0.08 | 0.00 | 4.5 | 4.6 |
| 2015 | 3166622_65:G>A | A10 | 15385404 | -0.08 | 0.00 | 5.1 | 7.4 |
| 2015 | 72478471:G>T | C04 | 3935715 | -0.07 | 0.00 | 3.0 | 5.0 |
| *Suggestive QTL* |  |  |  |  |  |  |  |
| 2015 | 3100298_18:T>C | A03 | 1903879 | -0.05 | 0.00 | 2.5 | 3.2 |
| 2014 | 4111998_10:T>C | A06 | 3099898 | 0.05 | 0.01 | 1.9 | 2.1 |
| 2015 | 30843166:A>G | A09 | 4927680 | 0.05 | 0.00 | 2.5 | 3.5 |
| 2014 | 4331930 | C03 | 1964260 | 0.10 | 0.01 | 2.2 | 6.9 |
| 2014 | 3091821_61:C>T | C08 | 20769516 | 0.05 | 0.01 | 2.0 | 2.2 |
| 2014 | 4706489 | C09 | 11162510 | 0.05 | 0.01 | 2.0 | 1.9 |

### QTL for NDVI

NDVI measurements were taken at periodically (7-14 days interval), starting from ∼6 weeks after sowing. All measurement taken during the course of experiments showed high correlation (r = 0.67 to 0.98) among genotypes suggesting that variation detected for NDVI is genetically determined (Supplemental Table 2). To identify and verify dynamic QTL for different NDVI measurements, we used all phenotypic scores taken before the onset of flowering. A total of 15 QTL (6 significant QTL with LOD ≥ 3 and nine suggestive QTLs with LOD score of <3) were detected on chromosomes A01, A02, A06, A07, A08, C02, C03 and C07 (Table 3). Three stable QTL on chromosomes A02, C02 and C03 were repeatedly detected in both environments. Of these, the QTL on chromosome C02 identified with a DArTseq marker 3092834 accounted for majority of the genetic variation in NDVI (R^2^ = 22.26%). Ag-Outback, the paternal parent of the DH population contributed alleles for higher NDVI on A02 and C02, whereas RP04 contributed alleles for higher NDVI at C03, suggesting that both parental alleles could enhance the fractional ground cover (early vigour) in canola.

**Table 3.** QTL associated with plant development and flowering time traits in a DH population from RP04/Ag-Outback. Loci that were repeatedly detected across environments are in highlighted in bold letters.

| Trait | Phenotyping year | Marker name | Chromosome | Physical position (bp) | Allelic effect | Probability | LOD | Genotypic variance explained (%) |
| --- | --- | --- | --- | --- | --- | --- | --- | --- |
| NDVI (14July) | 2015 | 4167794 | A01 | 10625625 | -0.01 | 0.00 | 2.77 | 9.13 |
| **NDVI (27July)** | **2015** | **3103028** | **A02** | **2609359** | **-0.02** | **0.00** | **3.72** | **12.50** |
| **NDVI(14July)** | **2015** | **3103028** | **A02** | **2609359** | **-0.02** | **0.00** | **4.52** | **15.20** |
| **NDVI (11Aug)** | **2015** | **3103028** | **A02** | **2609359** | **-0.02** | **0.00** | **3.31** | **11.06** |
| **NDVI (18July)** | **2014** | **3103028** | **A02** | **2609359** | **-0.02** | **0.00** | **4.00** | **13.46** |
| **NDVI (27June)** | **2014** | **3103028** | **A02** | **2609359** | **-0.02** | **0.00** | **3.08** | **10.25** |
| **NDVI (20June)** | **2014** | **3103028** | **A02** | **2609359** | **-0.02** | **0.00** | **3.08** | **10.23** |
| NDVI (18July) | 2014 | 3168358 | A02 | 6477720 | 0.04 | 0.00 | 3.62 | 12.16 |
| NDVI (27June) | 2014 | 3168358 | A02 | 6477720 | 0.03 | 0.00 | 2.42 | 7.83 |
| NDVI (20June) | 2014 | 3168358 | A02 | 6477720 | 0.03 | 0.00 | 2.66 | 8.71 |
| NDVI (27July) | 2015 | 3124936_62:A>C | A06 | 3982568 | -0.01 | 0.01 | 2.08 | 6.60 |
| NDVI (27June) | 2014 | 3102379_58:T>C | A07 | 8721875 | 0.02 | 0.00 | 2.59 | 8.45 |
| NDVI (20June) | 2014 | 3102379_58:T>C | A07 | 8721875 | 0.02 | 0.00 | 2.46 | 7.99 |
| NDVI (18July) | 2014 | 3139156_6:A>T | A08 | 18181217 | 0.01 | 0.00 | 3.00 | 9.94 |
| **NDVI (11Aug)** | **2015** | **100014951_59:T>C** | **C02** | **2526335** | **-0.02** | **0.00** | **3.92** | **13.19** |
| **NDVI (14July)** | **2015** | **3134538** | **C02** | **3021170** | **-0.03** | **0.00** | **6.48** | **21.43** |
| **NDVI (27July)** | **2015** | **3092834** | **C02** | **3718451** | **-0.04** | **0.00** | **6.76** | **22.26** |
| **NDVI (27June)** | **2014** | **3092834** | **C02** | **3718451** | **-0.03** | **0.00** | **4.49** | **15.13** |
| **NDVI (20June)** | **2014** | **3092834** | **C02** | **3718451** | **-0.02** | **0.00** | **4.10** | **13.80** |
| **NDVI (18July)** | **2014** | **3148811** | **C02** | **3781119** | **-0.03** | **0.00** | **6.12** | **20.35** |
| NDVI (27June) | 2014 | 7507904 | C02 | 7063310 | 0.02 | 0.00 | 2.96 | 9.79 |
| NDVI (20June) | 2014 | 7507904 | C02 | 7063310 | 0.02 | 0.00 | 2.70 | 8.87 |
| NDVI (18July) | 2014 | 3074777_66:C>A | C02 | 8042304 | 0.02 | 0.00 | 2.97 | 9.84 |
| NDVI (14July) | 2015 | 4335168 | C02 | 9110655 | 0.02 | 0.00 | 3.50 | 11.72 |
| NDVI (27July) | 2015 | 4113826 | C02 | 9375706 | 0.03 | 0.00 | 4.33 | 14.59 |
| NDVI (11Aug) | 2015 | 4113461_38:A>C | C02 | 9664537 | 0.03 | 0.00 | 4.02 | 13.53 |
| NDVI (27July) | 2015 | 3084367_42:A>T | C03 | 22640779 | 0.01 | 0.00 | 2.66 | 8.71 |
| NDVI (14July) | 2015 | 3084367_42:A>T | C03 | 22640779 | 0.01 | 0.00 | 3.63 | 12.17 |
| NDVI (11Aug) | 2015 | 3084367_42:A>T | C03 | 22640779 | 0.01 | 0.00 | 2.57 | 8.40 |
| **NDVI (14July)** | **2015** | **5810998** | **C03** | **23251374** | **0.01** | **0.00** | **2.37** | **7.65** |
| **NDVI (27June)** | **2014** | **5810998** | **C03** | **23251374** | **0.02** | **0.00** | **2.55** | **8.33** |
| **NDVI (20June)** | **2014** | **5810998** | **C03** | **23251374** | **0.02** | **0.00** | **2.56** | **8.34** |
| **NDVI (27July)** | **2015** | **5816633_59:G>A** | **C03** | **23287237** | **0.01** | **0.01** | **1.86** | **5.79** |
| NDVI (11Aug) | 2015 | 7251326 | C07 | 23499250 | 0.02 | 0.00 | 3.07 | 10.19 |
| NDVI (11Aug) | 2015 | 5053335_49:G>T | C07 | 33099868 | 0.02 | 0.00 | 2.73 | 8.96 |
| NDVI (27June) | 2014 | 5591686_47:G>C | C07 | 40274967 | 0.02 | 0.00 | 2.75 | 9.05 |
| NDVI (20June) | 2014 | 5591686_47:G>C | C07 | 40274967 | 0.02 | 0.00 | 2.72 | 8.93 |

### QTL for shoot biomass accumulation at vegetative stage

Six significant and five suggestive QTLs for shoot biomass were identified on chromosomes A01, A02, A05, A09, A10, C02, C03, C04, C05, C08 and C09 (Table 4). Of them, only one QTL on A01 delineated with 4119230_50:C>T and 5591791_60:A>T markers was repeatedly detected across both 2014 and 2015 environments, respectively and accounted up to 15.1% of the genotypic variance. The paternal parent, Ag-Outback contributed favourable allele for high biomass accumulation.

**Table 4.** QTL associated with shoot biomass accumulation in a DH population from RP04/Ag-Outback. Loci that were repeatedly detected across environments are in highlighted in bold letters.

| Trait | Phenotyping year | Marker name | Chromosome | Physical position (bp) | Allelic effect (g/plant) | Pr | LOD | Genotypic variance explained (%) |  |
| --- | --- | --- | --- | --- | --- | --- | --- | --- | --- |
| **BMFresh** | **2014** | **4119230_50:C>T** | **A01** | **21342507** | **-13.9** | **0.00** | **4.5** | **15.1** | **1** |
| **Biomass** | **2015** | **5591791_60:A>T** | **A01** | **21615163** | **-20.0** | **0.00** | **2.8** | **9.2** | **1** |
| Biomass | 2015 | 3097457 | A02 | 5951404 | 51.4 | 0.00 | 5.6 | 18.7 | 2 |
| BMFresh | 2014 | 3089785_23:T>C | A05 | 21615410 | -7.5 | 0.00 | 2.6 | 8.4 | 3 |
| BMFresh | 2014 | 4110682 | A09 | 33352074 | 12.2 | 0.00 | 4.9 | 16.6 | 4 |
| Biomass | 2015 | 4108375 | A10 | 4926878 | -20.9 | 0.00 | 3.7 | 12.6 | 5 |
| BMFresh | 2014 | 3148128_6:C>A | C02 | 36071167 | -8.6 | 0.00 | 2.8 | 9.4 | 6 |
| BMFresh | 2014 | 7507901 | C03 | 23251374 | 10.2 | 0.00 | 3.6 | 12.1 | 7 |
| BMFresh | 2014 | 3079920_42:A>G | C04 | 1567726 | 9.2 | 0.00 | 2.7 | 8.9 | 8 |
| BMFresh | 2014 | 3130517_15:A>G | C05 | 31690311 | 10.4 | 0.00 | 4.3 | 14.4 | 9 |
| BMFresh | 2014 | 3165286_42:T>C | C08 | 38018708 | -8.3 | 0.01 | 2.1 | 6.7 | 10 |
| Biomass | 2015 | 7250050_21:T>G | C09 | 800257 | -15.5 | 0.00 | 2.7 | 9.0 | 11 |

### QTL for flowering time

Flowering time was assessed as days to flowering at two stages: first flowering and the last flowering. Nine QTL (4 significant and 5 suggestive) were identified for first flowering on chromosomes A02, A05, A09, A10, C02, C03, C06 and C08 (Table 5). However, none of the QTL was repeatedly detected across environments. Maximum R^2^ (19.8%) was accounted by DArTseq marker; 3097457 on A02. For the last flowering time, six significant and three suggestive QTL were detected on chromosomes A01, A02, A05, A09, A10, C02, C06 and C08 (Table 5). These QTL accounted for 6.89% (A10) to 16.09% (C02) of the genotypic variance. Only one stable QTL on A05 was detected for 100% flowering time across environments, suggesting that G x E interactions play a role in revealing variation for flowering time. Both first and last flowering time showed high genetic correlation (*r^2^* > 0.9) within each year (Supplementary Table 1). This was also supported by our genetic analysis which revealed that ∼90% of QTLs (8/9) are mapped on the similar genomic positions of the reference Darmor-*bzh* genome (Table 5).

**Table 5.** QTL associated with shoot biomass accumulation in a DH population from RP04/Ag-Outback. Loci that were repeatedly detected across environments are in highlighted in bold letters.

| Trait | Phenotyping year | Marker name | Chromosome | Physical position (bp) | Allelic effect | Pr | LOD | Genotypic variance explained (%) |  |
| --- | --- | --- | --- | --- | --- | --- | --- | --- | --- |
| Last Flowering | 2015 | 3166097 | A01 | 3283242 | 1.1 | 0.00 | 3.3 | 11.0 | 1 |
| **First Flowering** | **2015** | **3097457** | **A02** | **5951404** | **-3.6** | **0.00** | **6.0** | **19.8** | **1** |
| **Last Flowering** | **2015** | **3097457** | **A02** | **5951404** | **-2.3** | **0.00** | **3.7** | **12.4** | **2** |
| **Last Flowering** | **2015** | **3085388_5:C>T** | **A02** | **23653368** | **1.1** | **0.00** | **3.6** | **12.0** | **3** |
| **First Flowering** | **2015** | **3150966** | **A02** | **24147662** | **1.1** | **0.00** | **2.5** | **8.1** | **2** |
| **First Flowering** | **2014** | **3139456** | **A05** | **1843494** | **-2.1** | **0.00** | **2.6** | **8.4** | **3** |
| **Last Flowering** | **2015** | **3139456** | **A05** | **1843494** | **1.0** | **0.00** | **3.1** | **10.4** | **4** |
| Last Flowering | 2014 | 5153681 | A05 | 2406544 | -1.6 | 0.01 | 2.3 | 7.4 | 4 |
| **First Flowering** | **2014** | **5818505_46:G>A** | **A09** | **5034799** | **1.6** | **0.02** | **1.6** | **4.9** | **4** |
| **Last Flowering** | **2014** | **5818505_46:G>A** | **A09** | **5034799** | **1.6** | **0.01** | **2.3** | **7.4** | **5** |
| **First Flowering** | **2014** | **3106140_65:T>G** | **A10** | **15487758** | **-1.9** | **0.00** | **2.3** | **7.5** | **5** |
| **Last Flowering** | **2014** | **3106140_65:T>G** | **A10** | **15487758** | **-1.5** | **0.01** | **2.2** | **6.9** | **6** |
| **First Flowering** | **2015** | **4112381** | **C02** | **2720764** | **1.7** | **0.00** | **4.2** | **14.2** | **6** |
| **Last Flowering** | **2015** | **4112381** | **C02** | **2720764** | **1.4** | **0.00** | **4.8** | **16.1** | **7** |
| First Flowering | 2014 | 3154725_22:A>G | C03 | 8035680 | 2.8 | 0.00 | 3.8 | 12.8 | 7 |
| **First Flowering** | **2014** | **5049884** | **C06** | **15906526** | **-2.4** | **0.00** | **3.3** | **11.2** | **8** |
| **Last Flowering** | **2014** | **5049884** | **C06** | **15906526** | **-1.6** | **0.00** | **2.5** | **8.2** | **8** |
| **Last Flowering** | **2015** | **3100323** | **C08** | **38039864** | **1.3** | **0.00** | **3.6** | **12.1** | **9** |
| **First Flowering** | **2015** | **3100323** | **C08** | **38063463** | **1.3** | **0.00** | **2.5** | **8.3** | **9** |

### Multi-trait QTL associations

Eight QTL associated with multiple trait were detected on A02, A07, A10, C2, C3, C4, C5 and C08 (Table 2, 3, 5, Figure 5). Of them, three DTI for seed yield related QTL were also associated with NDVI, flowering time and biomass; the first QTL region for NDVI and yield was detected on chromosome A06 within 0.9 Mb from each other (coordinates 3.09 Mb to 3.98 Mb), the second QTL region for flowering time, and DTI for seed yield was detected within coordinates 4.92 Mb to 5.03 Mb on chromosome A09, and the third QTL was detected on chromosome A10 for flowering time and DTI (15.39 Mb to 15.48 Mb (0.1 Mb interval encompassing several overlapping significant QTLs). One of the QTL for biomass and flowering time (First and last flowering) were collocated at the same position (coordinate 5951404) on chromosome A02 (Supplementary Table 2). These multiple trait associations suggest that these genomic regions for NDVI, flowering time and DTI are associated with correlated traits which may be either linked or represent to pleiotropic genes.

**Figure 5.**
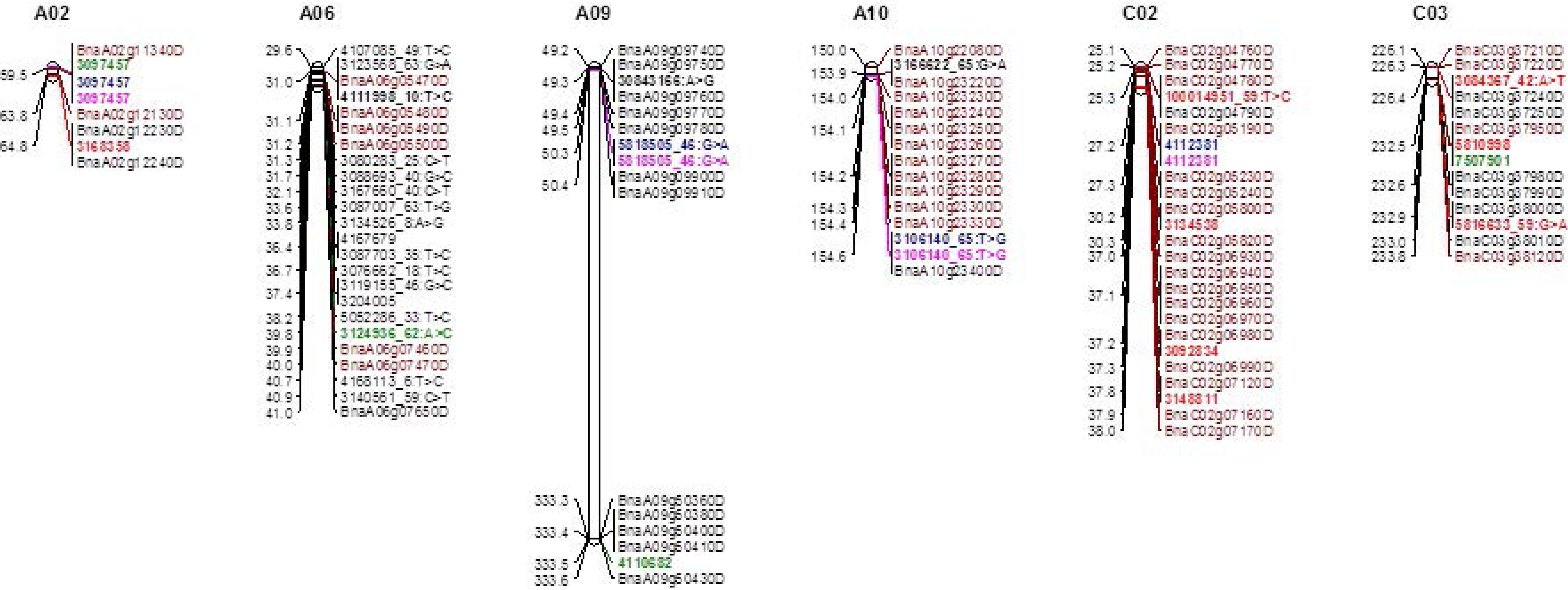
Partial linkage maps of the *B. napus* DH population derived from RP04/Ag-outback, showing the position of multi-trait QTL for normalised difference vegetative index (NDVI), shoot biomass, first flowering (25%), full flowering (100%) and seed yield. Markers which showed only significant trait-marker associations (LOD) are shown her in different colours: Red (NDVI), black (seed yield), green (shoot biomass), first flowering time (pink), full flowering time (blue) and candidate genes are highlighted in brown colour. Positions (in bp) of molecular markers and candidate genes are given on the Darmor assembly version 4; markers that were mapped on both genetic and physical maps are in bold and connected with solid lines. Graphical representations of linkage maps are made using MapChart. Linkage distances are given in cM (left side), while marker IDs are given in right hand side. Physical map distances are given in 1/100,000th fraction of the original coordinates for clarity.

### Physical mapping of significant trait-marker associations

To identify putative candidate genes underlying QTL implicated in trait variation, we searched the physical positions of markers that either appeared consistently across environments or were associated with multi-traits on the reference *B. napus* cv. Darmor-*bzh* genome assembly (http://www.genoscope.cns.fr/blat-server/cgi-bin/colza/webBlat). The list of candidate genes is presented in Table 6.

**Table 6:** Candidate genes located in the vicinity of significant QTL associations for different agronomic traits in a double haploid population from RP04/AG-Outback

| Trait | Phenotyping | Significant marker | Chromosome | Physical location of significant marker | *B. napus* annotated gene | Physical location of *B. napus* annotated gene | Distance (bp) between significant markers and the closest candidate gene | Annotated gene |
| --- | --- | --- | --- | --- | --- | --- | --- | --- |
| LAST FLOWERING | 2015 | 3166097 | A01 | 3283242 | BnaA01g06970D | 3282384 | 858 | Dihydropteroate synthase-like |
| Shoot biomass (g) | 2014 | 4119230_50:C>T | A01 | 21342507 | BnaA01g31300D | 21292552 | 61942 | DCD (Development and Cell Death) domain protein |
|  |  |  |  |  | BnaA01g31320D | 21280565 | 49955 | SAG20 (SENESCENCE ASSOCIATED GENE 20) |
| Shoot biomass (g) | 2015 | 5591791_60:A>T | A01 | 21615163 | BnaA01g31950D | 21606770 | 8393 | GASA5 (Gibberellin-regulated protein 5) |
|  |  |  |  |  | BnaA01g31960D | 21610805 | 4358 | Folylpolyglutamate synthetase, conserved site ATDFC-root development |
| NDVI (27th June) | 2014 | 3103028 | A02 | 2609359 | BnaA02g05670D | 2601299 | 8060 | alpha/beta-Hydrolases superfamily protein, ANAC090 (NAC domain-containing protein 90), |
| Shoot biomass (g) | 2015 | 3097457 | A02 | 5951404 | BnaA02g11340D | 5948180 | 3224 | Polycomb protein, VEFS-Box |
| NDVI (18th June) | 2014 | 3168358 | A02 | 6477720 | BnaA02g12230D | 6476887 | 833 | Late embryogenesis abundant protein, LEA-14 |
| Last flowering | 2015 | 3085388_5:C>T | A02 | 23653368 | BnaA02g33010D | 23649506 | 3862 | ENTH/VHS |
| First flowering | 2014 | 3139456 | A05 | 1843494 | BnaA05g03250D | 1815458 | 28036 | F-box/kelch-repeat protein (F-box associated interaction domain) |
| Biomass | 2014 | 3089785_23:T>C | A05 | 21615410 | BnaA05g31350D | 21616785 | 1375 | Major facilitator superfamily domain |
| Yield (t/ha) | 2014 | 31312620:A>G | A06 | 19833237 | BnaA06g29020D | 19833237 | 1897 | Seed maturation protein |
| NDVI (18th July) | 2014 | 3139156_6:A>T | A08 | 18181217 | BnaA08g27140D | 18179574 | 1643 | Cytochrome c oxidase assembly protein CtaG/Cox11, domain |
|  |  |  |  |  | BnaA08g27150D | 18181524 | 307 | Pentatricopeptide repeat |
| Biomass | 2014 | 4110682 | A09 | 33352074 | BnaA09g50430D | 33356001 | 3927 | Lipase/lipoxygenase, PLAT/LH2 |
| Biomass | 2015 | 4108375 | A10 | 4926878 | BnaA10g06460D | 4915483 | 11295 | Zinc finger, RING/FYVE/PHD-type |
|  |  |  |  |  | BnaA10g06470D | 4928841 | 1963 | Spermidine/spermine synthases family |
| Yield (t/ha) | 2014 | 3166622_65:G>A | A10 | 15398506 | BnaA10g23220D | 15394043 |  | Zinc finger, RING/FYVE/PHD-type |
| NDVI (11th Aug) | 2015 | 100014951_59:T>C | C02 | 2526335 | BnaC02g04790D | 2526638 | 303 | G-protein beta WD-40 repeat |
| First flowering | 2015 | 4112381 | C02 | 2720764 | BnaC02g05190D | 2718285 | 2479 | Ankyrin repeat-containing domain |
| NDVI (14th July) | 2015 | 3134538 | C02 | 3021170 | BnaC02g05800D | 3020889 | 281 | NHL repeat, subgroup |
| NDVI ( 27th July) | 2015 | 3092834 | C02 | 3718451 | BnaC02g06930D | 3698436 | 20015 | AtCslD2 (Cellulose synthase-like protein D2) |
| NDVI (18th July) | 2014 | 3148811 | C02 | 3781119 | BnaC02g07130D | 3782444 | 1325 | Unknown protein |
|  |  |  |  |  | BnaC02g07170D | 3795126 | 14007 | Rubisco LS methyltransferase, substrate-binding domain |
| NDVI (27th June) | 2014 | 7507904 | C02 | 7063310 | BnaC02g11790D | 7058173 | 5137 | Phosphoesterase At2g46880 |
| NDVI (18th July) | 2014 | 3074777_66:C>A | C02 | 8042304 | BnaC02g12590D | 8036959 | 5345 | Leucine rich repeat 4 |
| NDVI (14th July) | 2015 | 4335168 | C02 | 9110655 | BnaC02g13780D | 9109883 | 772 | Unknown protein |
|  |  |  |  |  | BnaC02g13810D | 9127855 | 17200 | Stomatin |
| NDVI (27July) | 2015 | 4113826 | C02 | 9375706 | BnaC02g14110D | 9378151 | 2445 | Cystathionine beta-synthase, core |
| NDVI (11Aug) | 2015 | 4113461_38:A>C | C02 | 9664537 | BnaC02g14280D | 9663793 | 744 | Protein of unknown function DUF295 |
| FIRST FLOWERING | 2014 | 3154725_22:A>G | C03 | 8035680 | BnaC03g15910D | 8029356 | 6324 | Armadillo-type fold |
| NDVI (14th July) | 2015 | 3084367_42:A>T | C03 | 22640779 | BnaC03g37240D | 22640798 | 19 | Uncharacterised protein family UPF0497, trans-membrane plant |
| NDVI (14th July) | 2015 | 5810998 | C03 | 23251374 | BnaC03g37950D | 23249334 | 2040 | ECT2 ( evolutionarily conserved C-terminal region 2, YTH domain) |
| Yield (t/ha) | 2014 | 72478471:G>T | C04 | 3935715 | BnaC04g05540D | 3935485 | 230 | F-box and associated interaction domain |
| Shoot biomass (g) | 2014 | 3130517_15:A>G | C05 | 31690311 | BnaC05g32070D | 31495290 | 195021 | F-box associated interaction domain |
|  |  |  |  |  | BnaC05g32060D | 31479559 | 210752 | Cytochrome P450 |
|  |  |  |  |  | BnaC05g32100D | 31540163 | 150148 | Spermidine/spermine synthases family |
|  |  |  |  |  | BnaC05g32120D | 31776235 | 85924 | Photosystem II PsbC |
| FIRST FLOWERING | 2014 | 5049884 | C06 | 15906526 | BnaC06g13140D | 15916851 | 10325 | Unknown |
| NDVI (11Aug) | 2015 | 7251326 | C07 | 23499250 | BnaC07g17320D | 23463976 | 35274 | ATP-sulfurylase PUA-like domain |
| Last flowering | 2015 | 3100323 | C08 | 38039864 | BnaC08g45320D | 38040135 | 271 | RABA2b (Ras-related protein Small GTPase superfamily, ARF type) |
|  |  |  |  |  | BnaC08g45310D | 38037731 | 2133 | Leucine rich repeat 4 |

Significant QTL for DTI in terms of seed yield were located within 5 kb from seed maturation protein, Zinc finger, RING/FYVE/PHD-type and F-box and associated interaction domain. QTL for flowering time were located within 30kb from F-box associated interaction domain, and Putative S-adenosyl-L-methionine-dependent methyltransferase (BnaC03g15890D). For NDVI, QTL were located near several genes of interest such as NAC domain-containing protein 90; ANAC090 (BnaA02g05600D), *FLOWERING LOCUS T* (BnaA02g12130D, BnaA07g132090D), DELLA (BnaA02g12260D), lateral organ (BnaA02g12180D), Late embryogenesis abundant protein (LEA-14;BnaA02g12230D), Cellulose synthase-like protein D2 (BnaC02g06930D), Fasciclin-like arabinogalactan family protein (BnaC02g06940D), and Cytochrome c oxidase (BnaA08g27140D, BnaC02g11870D). QTL for shoot biomass were located within 0.2 Mb from DEVELOPMENT/CELL DEATH, Gibberellin-regulated protein 5, senescence-associated gene 20, flowering time genes (polycomb group protein), RING-H2 finger protein 2B, F-box associated interaction genes.

## Discussion

Genetic improvement in seed yield and its stability under water-limited conditions is one of the major objectives of canola breeding programs. Improving drought tolerance by selecting genotypes based on their response to drought stress on yield and productivity traits such as biomass accumulation, flowering time, and water-use efficiency will help to develop new varieties with relatively stable seed yield across different environments. In canola, and its close relatives *B. rapa* and *B. oleracea*, only a limited number of component traits such as leaf weight, stomatal conductance, carbon isotope discrimination, and root pulling force [19, 20, 29] have been investigated to gain an understanding on the genetic basis of drought tolerance.

Australia is the driest continent on earth and crop productivity heavily relies on in-season rain and/or on the stored moisture. Canola breeding programs have been selecting varieties for high yield under Australian conditions from last 40 years and may have selected varieties for improved productivity unintentionally without prior knowledge of loci associated with resilience to drought stress. Genetic loci associated with yield across a range of low average rainfall environments (drought tolerance) have not been identified in Australian canola germplasm as yet. Herein, we implemented an incremental crop tolerance approach to identify genomic regions associated with response to terminal drought stress for seed yield. We identified twenty five QTL controlling genetic variation in plasticity traits in a DH population from RP04/Ag-Outback (Table 2, 3, 5). Of them, at least eight multi-trait QTL for flowering time, NDVI, shoot biomass and seed yield were identified on chromosomes A02, A07, A10, C02, C03, C04, and C08 (Figure 5, Supplementary Table 2). This observed clustering of different QTL is consistent with genetic correlation among different traits in the RADH population (Supplementary Table 1). In a previous study, Raman et al [30] found moderate to high genetic correlations for flowering time, early vigour, plant biomass and seed yield in an Australian DH population from Skipton/Ag-Spectrum//Skipton. Flowering time is an adaptive trait and shows pleiotropy with seed yield and other plant architectural traits in *B. juncea* and winter/spring crosses of *B. napus* [19, 23, 30–34]. However, in the present study, we used Australian spring type genotypes to minimise the effect of vernalisation genes on flowering time and seed yield.

We attempted to locate the candidate genes within 1Mbp from significant trait-marker associations; however these genes may not be causative for trait variations and require further validation studies. Nevertheless, these genomic regions provide evidence that QTL likely represent to ‘regions of interest’. In addition, a *priori* candidate genes identified in the vicinity of QTL in this study have been shown to control trait variation in other species, suggesting that the trait-marker associations identified herein are valid. For examples, some of the QTL representing trait(s) are located near the flowering time genes such as *FLOWERING LOCUS T, Polycomb Response Complex 2 (BnaA02g11340D)*, SAM (BnaA07g11700D), FLOWERING LOCUS C (BnaA10g22080D), *PHYC*, *TEM1*, GIGENTITA (GI), and *POWERDRESS* (Figure 6). These results are consistent with a previous study, which reported a multi-trait QTL (flanked by markers 3110489 and 3075574) for plant emergence, shoot biomass, flowering time, and seed yield mapped on chromosome A07 in the vicinity of the *FT* [30]. PRC2 is involved in trimethylation of histone H3 lysine 27 and regulates VERNALISATION INSENSTIVE 3 (VIN3) and plays an important role in epigenetic repression of FLC [35]. GI is known to accelerate flowering under water stress by promoting *FT* and *TWIN SISTERS of FT* (*TSF*) genes via ABA signalling [36]. VIN3 is also implicated in adaptation to low oxygen conditions [37]. Likewise, phytochromes; especially *PHYB* are also implicated in increasing drought tolerance by enhancing ABA sensitivity when soil water becomes a limiting factor for sustaining plant growth [38]. One of the suggestive multi-trait QTL regions underling DTI and flowering time on A09 was located within 4 kb from Gibberellin regulated protein (BnaA09g09740D) and F-box associated interaction domain (BnaA09g09900D). This was further supported by our results on localisation of QTL for seed yield and flowering time (chromosome A10) near the *BnFLC.A10* gene involved in vernalisation response, Bna.ATE1 (BnaA10g24850D) implicated in germination and early vigour [39], Cytochrome P450, and HUA2-like 1 genes involved in upregulation of *FLC* [40, 41]. Hatzig et al [42] also identified significant association for elongation speed of radicle in *B. napus* GWAS panel of 248 lines within the marker interval on chromosome A10 Our findings are also consistent with a previous study which reported a close link between flowering time on *BnFLC.A10* and response to drought stress in a DH population derived from IMC106RR/Wichita [19, 43]. Therefore, it is likely that flowering time genes may be driving phenotypic plasticity to water stress in the RADH population. We located QTL for NDVI near the GA_3_ and related genes such as DELLA and putative F-box protein which positively regulate GA signalling (Wood et al 2006, de Lucia et al 2008). Gibberellins are known in differentiation and elongation of cell/growth. Genes encoding Zinc finger, Ring type/PHD type proteins are implicated in seed development and yield [44, 45] and folypolyglutamate synthtase in primary root development [46]. Alpha/beta-Hydrolases proteins are involved in the formation of cutin, a waxy substance, which combine with cellulose forms a substance nearly impervious to water and constituting the cuticle in plants [47]. Detection of candidates hint that the described genes are the likely targets to gain further understanding on their functions and gene-networks involved in drought tolerance in canola. Taken together, the QTL regions identified herein may be involved in drought avoidance and/or drought tolerance mechanisms in canola.

Our results showed that DTI for seed yield is a complex trait controlled by several QTL and environment interaction (Table 2). However other correlated traits that are simple to measure such as NDVI could be used as a proxy to select drought tolerant genotypes in canola under water-limited conditions. Our results showed that NDVI correlates well with seed yield (Figure 2), suggesting that genetic improvement in seed yield can be made by selecting highly vigorous genotypes in canola breeding programs. Although in this study, we did not investigate the link between NDVI and seed yield under contrasting water regimes. Under water-limited conditions, early flowering/maturing varieties may yield higher compared to late flowering one, as plants trade-off seed yield with plant biomass. This scenario of drought escape mechanism often occurs in terminal drought under the Mediterranean environments prevalent in Australia and elsewhere. Varieties having early vigour are likely to have better germination rate, seedling vigour and better ground cover, saving available valuable resources such as water and nutrients; which may lead to early light inception and plant growth, enabling plants to flower early. Under terminal drought environments, flowering time and seed yield generally showed negative correlation; however the degree of correlation varied across environments [19, 23]. It was interesting to find no correlation (−0.06) between flowering time and DTI for seed yield in present study across two environment (Figure 3), suggesting that parental lines may have plasticity for drought escape (early flowering) as well as drought tolerance (water use efficiency). G × E can play an important role in dissection of trait-marker association. For example, in *B. rapa*, 54 QTL associated with 20 physiological and morphological traits involved in plant performance under control and drought conditions were identified. Of which, 17 QTL showed significant QTL-environment interactions [20]. Under irrigated conditions, both group of lines: (i) high vigour and early flowering, and (ii) low vigour and late flowering varieties may not correlate well due to the delay in flowering time and longer pod/seed filling window, compared to under terminal drought situations. It is also possible that late flowering lines may have genes for improved water-use efficiency/water productivity, therefore contributing to drought tolerance mechanism. Further research is required to validate this hypothesis.

## Conclusions

In summary, this study employed the incremental crop tolerance approach to determine the genetic variation in drought tolerance and to identify QTL for drought tolerance index in canola. We identified 25 QTL for NDVI, above ground shoot biomass at vegetative stage, flowering time, and seed yield in response to drought stress; majority of them had small allelic effects in DH lines. Both parental lines carried favourable alleles for enhancing adaptation under drought. Several putative candidate genes, such as *BnFLC.A10* and *Bna.ATE1* were localised within genomic regions associated with plasticity traits in response to drought stress. QTL regions were delimited with genotyping by sequencing based markers, which could be deployed to capture favourable alleles in the canola breeding program using marker-assisted/genomic selection strategies. We anticipate that improved selection of the plasticity traits can lead to increase in genetic gain for drought tolerance thus improving canola yield in drier environments. Further research is required to test the utility of the QTL identified in this study for improving canola yield under drought conditions across canola production regions.

## Data Availability

The seeds of the accessions are available from the Victorian Department of Agriculture, Horsham, Australia

## Supporting Information

Supplementary Figure 1: Meteorological data of phenotypic environments in 2014 and 2015

Supplementary Figure 2: Frequency histogram showing the distribution of genotypic means of DH lines from RP04/Ag-Outback.

Supplementary Table 1: Genetic correlation between traits evaluated across two consecutive years (2014 and 2015) in the DH population from RP04/Ag-Outback.

Supplementary Table 2: Multi trait QTL identified in the DH population from RP04/Ag-Outback.

## Acknowledgments

The authors would like to thank Dr Brian Cullis, University of Wollongong for discussion and guidance in modelling the incremental crop tolerance index, Mr David Leah, Mr Dave Roberts and Chris Fuller for sowing field trials and NDVI measurement.

