## Supplemental Figure for "QTL mapping reveals genomic regions for yield based on incremental tolerance index to drought stress and related agronomic traits in canola"

Supplementary Figure 1: Monthly rainfall during growing season at Wagga Wagga in 2014 and 2015

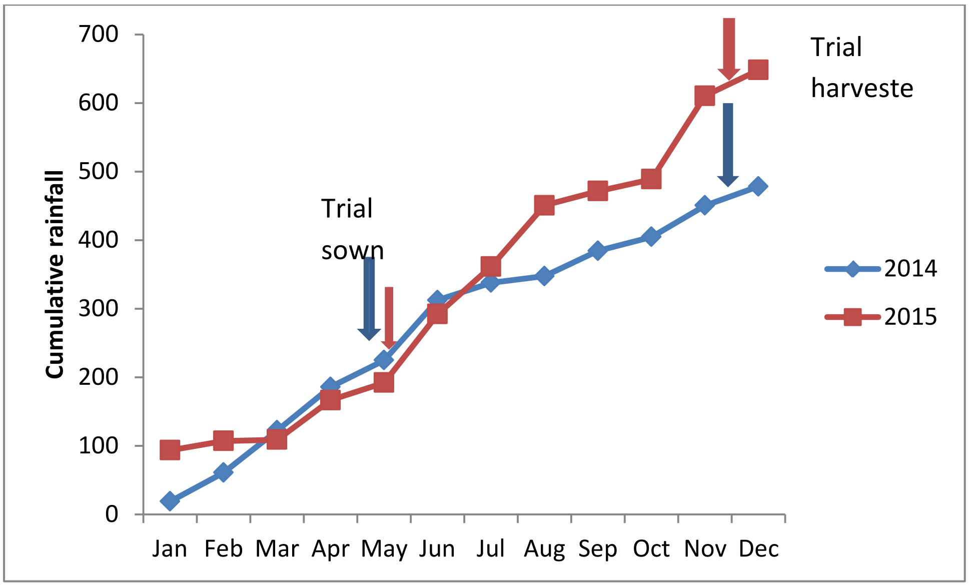

Supplementary Figure 2: Frequency histogram showing the distribution of genotypic means of DH lines from RP04/Ag-Outback for drought tolerance index (A-D), flowering time (E-F), shoot biomass (G-H), and NDVI (I-J). Parental means of the RP04 (RP)/Ag-Outback (AG) derived DH populations are shown with down arrows.

A

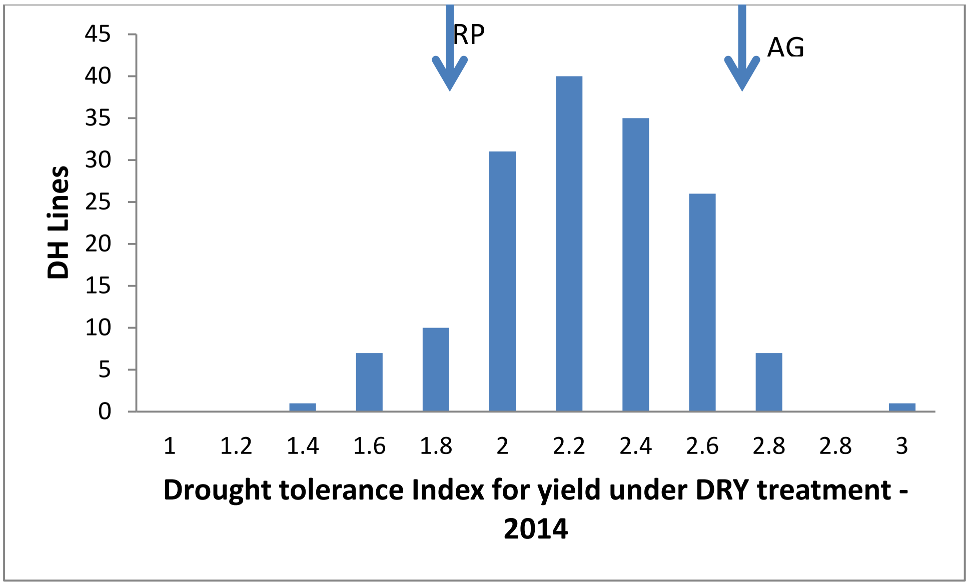

B

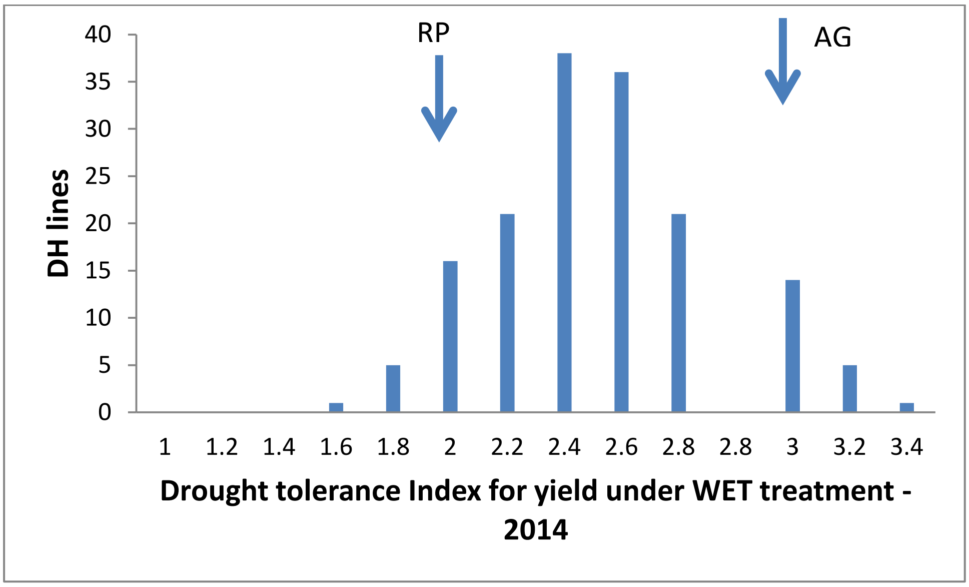

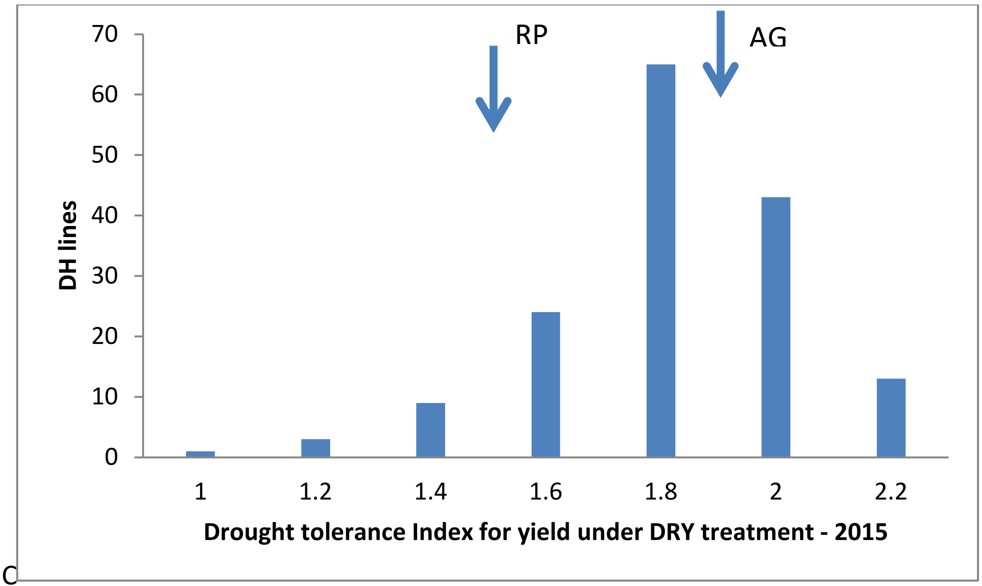
RP
AG
70
60
50
40
30
20
10
0
DH lines
1
1.2
1.4
1.6
1.8
2
2.2
Drought tolerance Index for yield under DRY treatment - 2015

C

D

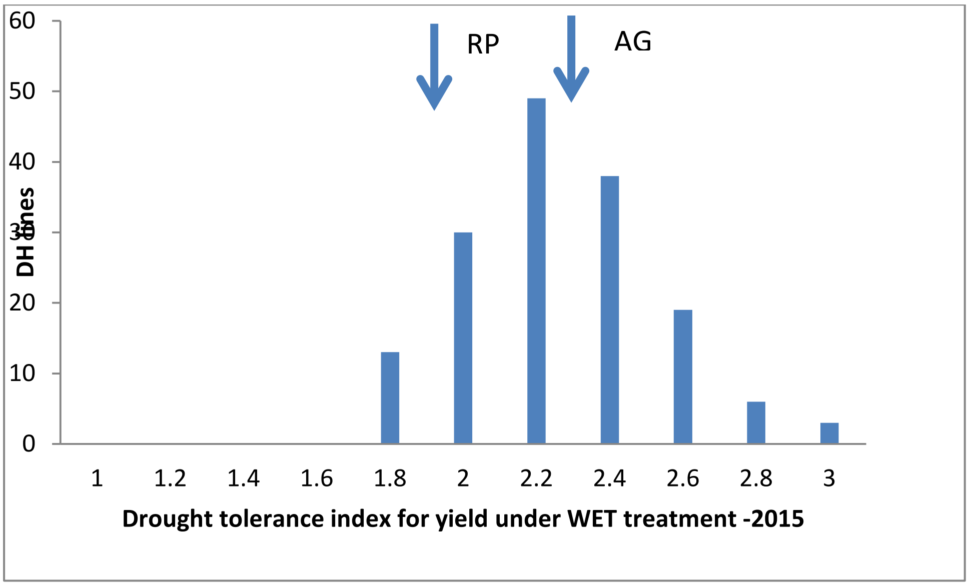
RP
AG
60
50
40
30
20
10
0
DH lines
1
1.2
1.4
1.6
1.8
2
2.2
2.4
2.6
2.8
3
Drought tolerance index for yield under WET treatment -2015

E

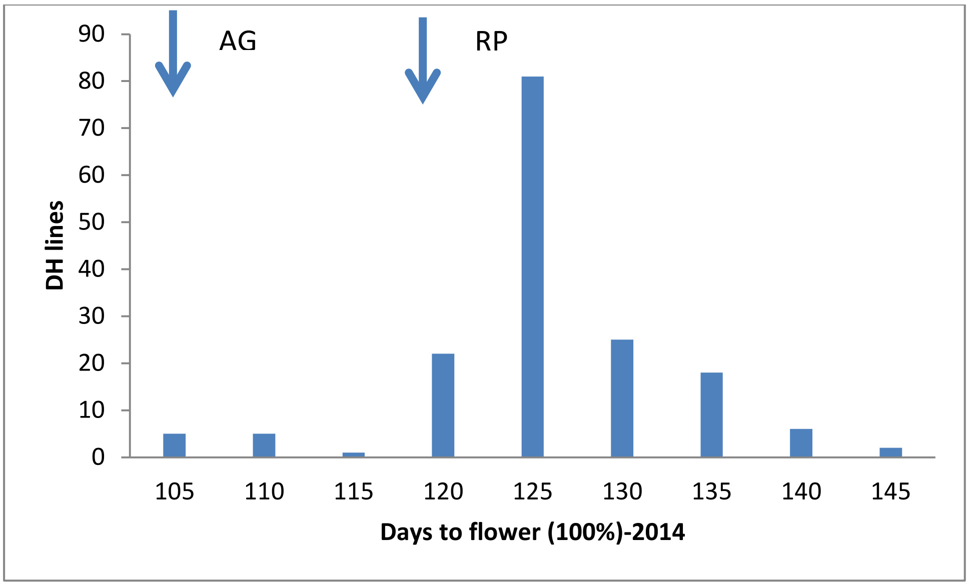
AG
RP
DH lines
90
80
70
60
50
40
30
20
10
0
105
110
115
120
125
130
135
140
145
Days to flower (100%)-2014

F

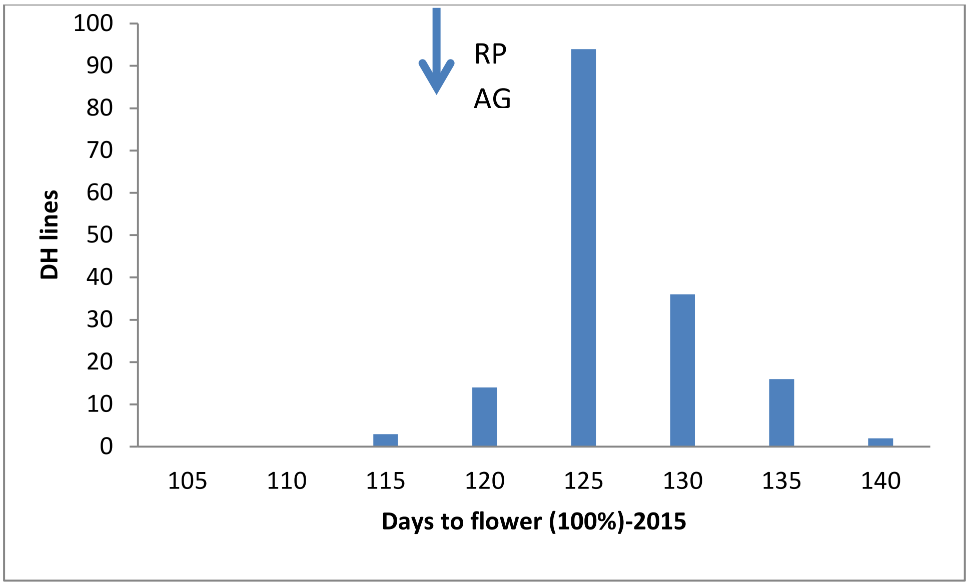
RP
AG
DH lines
100
90
80
70
60
50
40
30
20
10
0
105
110
115
120
125
130
135
140
Days to flower (100%)-2015

G

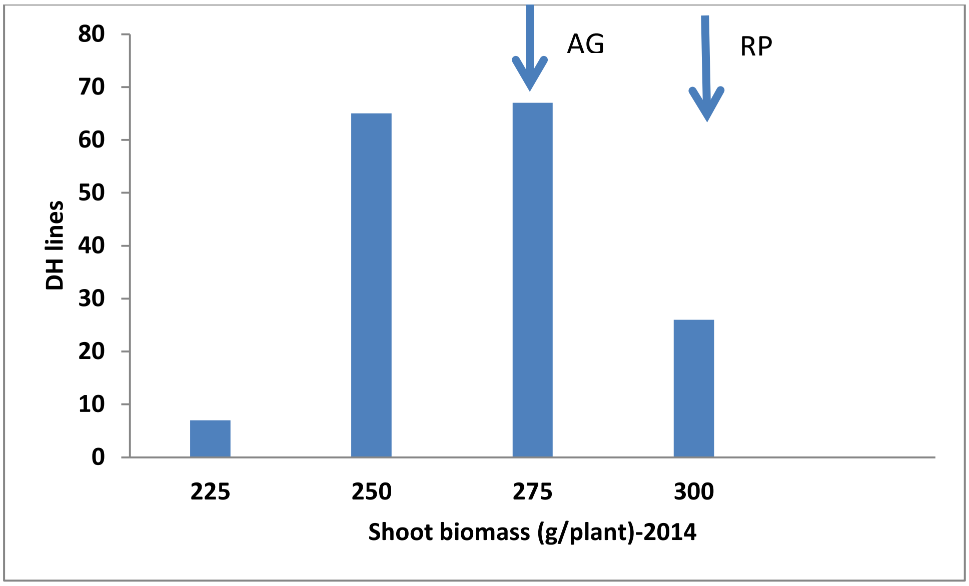
AG
RP
80
70
60
50
40
30
20
10
0
DH lines
225
250
275
300
Shoot biomass (g/plant)-2014

H

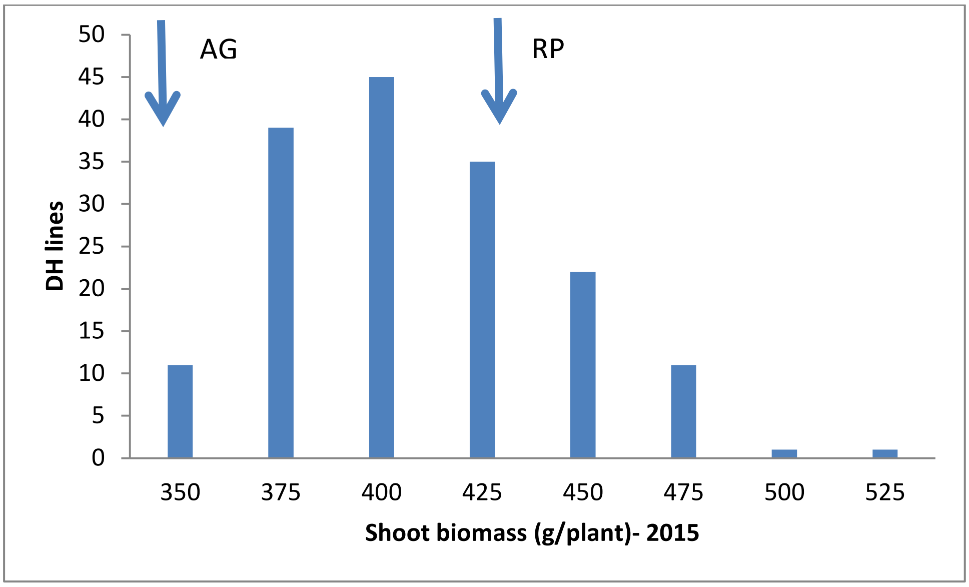
AG
RP
50
45
40
35
30
25
20
15
10
5
0
DH lines
350
375
400
425
450
475
500
525
Shoot biomass (g/plant)- 2015

I

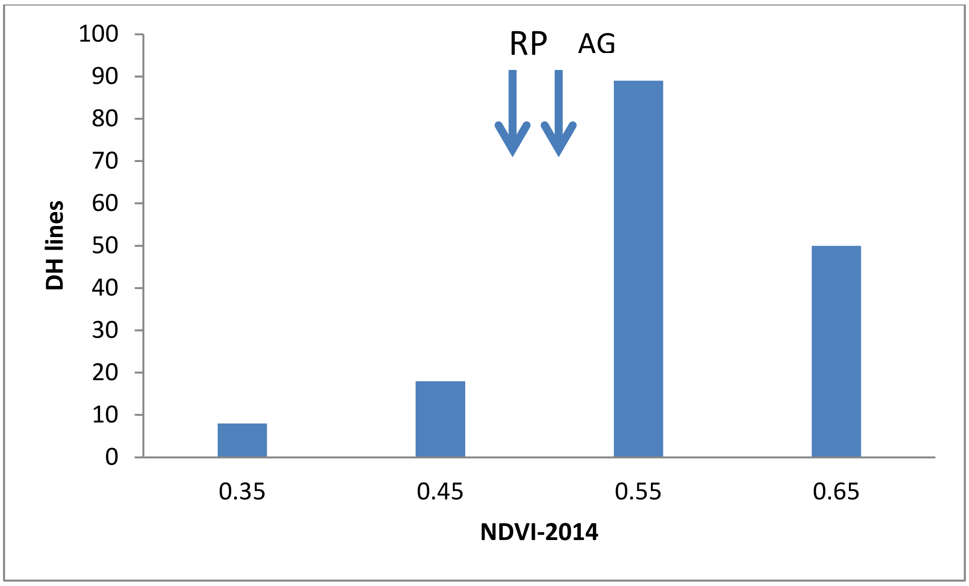
RP
AG
100
90
80
70
60
50
40
30
20
10
0
DH lines
0.35
0.45
0.55
0.65
NDVI-2014

J

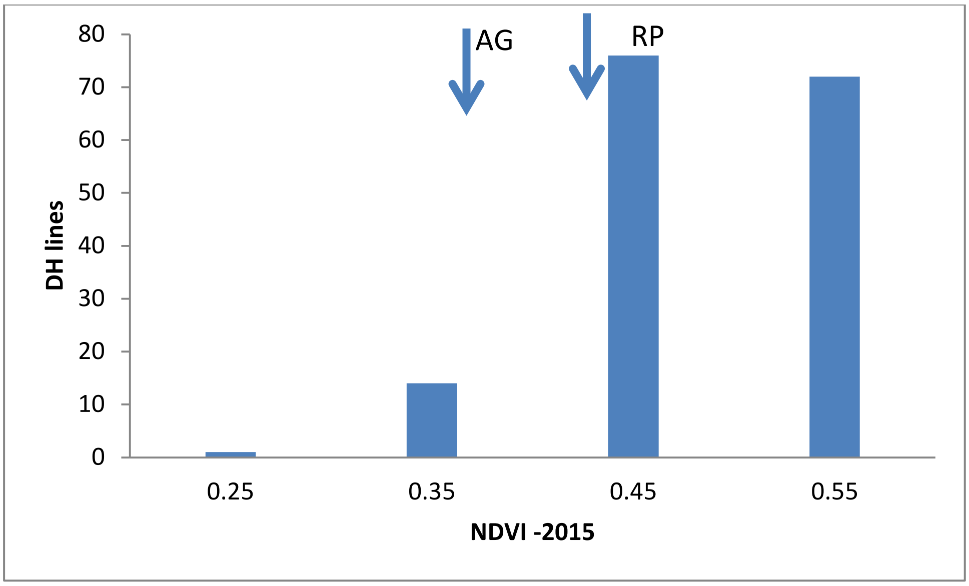
AG
RP
80
70
60
50
40
30
20
10
0
DH lines
0.25
0.35
0.45
0.55
NDVI -2015

**Supplementary Table 1: Genetic correlation between traits evaluated under WET**

| | | Biomass (g | Eabslishme | Ful (100%) | First flowe | Flowering ( | Plant heigh |
|---|---|---|---|---|---|---|---|
| | | | | | | | 2014 |
| 2014 | Biomass (g/plant) | 1.000 | 0.048 | -0.058 | -0.034 | -0.021 | 0.152 |
| 2014 | Eabslishment score (1-9) | 0.048 | 1.000 | -0.089 | -0.071 | 0.044 | 0.054 |
| 2014 | Ful (100%) flowering | -0.058 | -0.089 | 1.000 | 0.920 | -0.342 | 0.111 |
| 2014 | First flowering (25%) | -0.034 | -0.071 | 0.920 | 1.000 | -0.659 | 0.105 |
| 2014 | Flowering duration (days) | -0.021 | 0.044 | -0.342 | -0.659 | 1.000 | -0.052 |
| 2014 | Plant height (cm) | 0.152 | 0.054 | 0.111 | 0.105 | -0.052 | 1.000 |
| 2014 | NDVI (20Jun14) | 0.268 | 0.755 | -0.079 | -0.065 | 0.034 | 0.235 |
| 2014 | NDVI (27Jun14) | 0.271 | 0.762 | -0.080 | -0.070 | 0.038 | 0.241 |
| 2014 | NDVI (8Jul14) | 0.224 | 0.746 | -0.075 | -0.079 | 0.070 | 0.227 |
| 2014 | Grain yield(T/ha) | 0.136 | 0.190 | -0.103 | -0.069 | -0.033 | 0.514 |
| 2015 | Biomass (g/plant) | 0.380 | 0.146 | -0.029 | -0.047 | 0.067 | 0.204 |
| 2015 | Eabslishment score (1-9) | 0.153 | 0.431 | -0.142 | -0.053 | -0.099 | 0.007 |
| 2015 | Ful (100%) flowering | -0.264 | -0.339 | 0.151 | 0.111 | -0.018 | 0.299 |
| 2015 | First flowering (25%) | -0.258 | -0.358 | 0.176 | 0.141 | -0.047 | 0.321 |
| 2015 | Flowering duration (days) | 0.171 | 0.241 | -0.199 | -0.186 | 0.107 | -0.229 |
| 2015 | Plant height (cm) | 0.107 | -0.019 | 0.134 | 0.099 | 0.004 | 0.889 |
| 2015 | NDVI (11Aug15) | 0.244 | 0.420 | -0.057 | 0.000 | -0.071 | 0.417 |
| 2015 | NDVI (14Jul15) | 0.295 | 0.534 | -0.061 | -0.016 | -0.049 | 0.268 |
| 2015 | NDVI (27Jul15) | 0.283 | 0.518 | -0.036 | 0.020 | -0.085 | 0.300 |
| 2015 | Plant density (plant no./1 li | 0.021 | 0.352 | -0.088 | -0.021 | -0.096 | 0.115 |
| 2015 | Grain yield (t/Ha) | 0.033 | 0.115 | -0.063 | -0.010 | -0.098 | 0.422 |

¨ and DRY conditions across two consecutive years.

| NDVI (20Ju | NDVI (27Ju | NDVI (8Jul | Grain yield | Biomass (g | Eabslishme | Ful (100%) | First flowe | Flowering ( |
|---|---|---|---|---|---|---|---|---|
| 0.268 | 0.271 | 0.224 | 0.136 | 0.380 | 0.153 | -0.264 | -0.258 | 0.171 |
| 0.755 | 0.762 | 0.746 | 0.190 | 0.146 | 0.431 | -0.339 | -0.358 | 0.241 |
| -0.079 | -0.080 | -0.075 | -0.103 | -0.029 | -0.142 | 0.151 | 0.176 | -0.199 |
| -0.065 | -0.070 | -0.079 | -0.069 | -0.047 | -0.053 | 0.111 | 0.141 | -0.186 |
| 0.034 | 0.038 | 0.070 | -0.033 | 0.067 | -0.099 | -0.018 | -0.047 | 0.107 |
| 0.235 | 0.241 | 0.227 | 0.514 | 0.204 | 0.007 | 0.299 | 0.321 | -0.229 |
| 1.000 | 0.979 | 0.894 | 0.413 | 0.308 | 0.534 | -0.392 | -0.421 | 0.314 |
| 0.979 | 1.000 | 0.930 | 0.432 | 0.302 | 0.527 | -0.392 | -0.432 | 0.342 |
| 0.894 | 0.930 | 1.000 | 0.399 | 0.295 | 0.510 | -0.350 | -0.401 | 0.343 |
| 0.413 | 0.432 | 0.399 | 1.000 | 0.204 | 0.269 | -0.063 | -0.075 | 0.073 |
| 0.308 | 0.302 | 0.295 | 0.204 | 1.000 | 0.100 | -0.206 | -0.263 | 0.372 |
| 0.534 | 0.527 | 0.510 | 0.269 | 0.100 | 1.000 | -0.335 | -0.369 | 0.277 |
| -0.392 | -0.392 | -0.350 | -0.063 | -0.206 | -0.335 | 1.000 | 0.964 | -0.491 |
| -0.421 | -0.432 | -0.401 | -0.075 | -0.263 | -0.369 | 0.964 | 1.000 | -0.691 |
| 0.314 | 0.342 | 0.343 | 0.073 | 0.372 | 0.277 | -0.491 | -0.691 | 1.000 |
| 0.156 | 0.169 | 0.167 | 0.417 | 0.201 | -0.014 | 0.341 | 0.348 | -0.191 |
| 0.670 | 0.683 | 0.678 | 0.508 | 0.298 | 0.686 | -0.074 | -0.133 | 0.219 |
| 0.772 | 0.765 | 0.707 | 0.428 | 0.302 | 0.737 | -0.315 | -0.351 | 0.286 |
| 0.747 | 0.753 | 0.720 | 0.444 | 0.309 | 0.743 | -0.242 | -0.293 | 0.285 |
| 0.447 | 0.422 | 0.393 | 0.312 | -0.006 | 0.717 | -0.143 | -0.165 | 0.125 |
| 0.319 | 0.348 | 0.357 | 0.733 | 0.106 | 0.342 | 0.062 | 0.044 | -0.010 |

| Plant heigh | NDVI (11A | NDVI (14Ju | NDVI (27Ju | Plant dens | Grain yield (t/Ha) |
|---|---|---|---|---|---|
| 2015 | | | | | |
| 0.107 | 0.244 | 0.295 | 0.283 | 0.021 | 0.033 |
| -0.019 | 0.420 | 0.534 | 0.518 | 0.352 | 0.115 |
| 0.134 | -0.057 | -0.061 | -0.036 | -0.088 | -0.063 |
| 0.099 | 0.000 | -0.016 | 0.020 | -0.021 | -0.010 |
| 0.004 | -0.071 | -0.049 | -0.085 | -0.096 | -0.098 |
| 0.889 | 0.417 | 0.268 | 0.300 | 0.115 | 0.422 |
| 0.156 | 0.670 | 0.772 | 0.747 | 0.447 | 0.319 |
| 0.169 | 0.683 | 0.765 | 0.753 | 0.422 | 0.348 |
| 0.167 | 0.678 | 0.707 | 0.720 | 0.393 | 0.357 |
| 0.417 | 0.508 | 0.428 | 0.444 | 0.312 | 0.733 |
| 0.201 | 0.298 | 0.302 | 0.309 | -0.006 | 0.106 |
| -0.014 | 0.686 | 0.737 | 0.743 | 0.717 | 0.342 |
| 0.341 | -0.074 | -0.315 | -0.242 | -0.143 | 0.062 |
| 0.348 | -0.133 | -0.351 | -0.293 | -0.165 | 0.044 |
| -0.191 | 0.219 | 0.286 | 0.285 | 0.125 | -0.010 |
| 1.000 | 0.382 | 0.223 | 0.262 | 0.084 | 0.395 |
| 0.382 | 1.000 | 0.847 | 0.903 | 0.571 | 0.536 |
| 0.223 | 0.847 | 1.000 | 0.951 | 0.583 | 0.370 |
| 0.262 | 0.903 | 0.951 | 1.000 | 0.581 | 0.427 |
| 0.084 | 0.571 | 0.583 | 0.581 | 1.000 | 0.281 |
| 0.395 | 0.536 | 0.370 | 0.427 | 0.281 | 1.000 |

Supplementary Table 2: Multi trait QTL identified in the DH population from RP04/Ag-Outback.

| Trait | Phenotyping year | Marker name | Chromosome | Physical position (bp) | Allelic effect (t/ha) | Probability (P =0.05) |
|---|---|---|---|---|---|---|
| Biomass | 2015 | 3097457 | A02 | 5951404 | 51.40413 | 2.53E-06 |
| FIRST FLOWERING | 2015 | 3097457 | A02 | 5951404 | -3.62691 | 1.1E-06 |
| LAST FLOWERING | 2015 | 3097457 | A02 | 5951404 | -2.27017 | 0.000205 |
| DTI (seed yield) | 2014 | 4111998_10:T>C | A06 | 3099898 | 0.05 | 0.01 |
| NDVI (27July) | 2015 | 3124936_62:A>C | A06 | 3982568 | -0.01 | 0.01 |
| DTI (seed yield) | 2015 | 30843166:A>G | A09 | 4927680 | 0.05 | 0 |
| FIRST FLOWERING | 2014 | 5818505_46:G>A | A09 | 5034799 | 1.644139 | 0.024256 |
| LAST FLOWERING | 2014 | 5818505_46:G>A | A09 | 5034799 | 1.640575 | 0.005143 |
| DTI (seed yield) | 2015 | 3166622_65:G>A | A10 | 15385404 | -0.08 | 0 |
| FIRST FLOWERING | 2014 | 3106140_65:T>G | A10 | 15487758 | -1.88866 | 0.004719 |
| LAST FLOWERING | 2014 | 3106140_65:T>G | A10 | 15487758 | -1.45263 | 0.006892 |
| NDVI (11Aug) | 2015 | 100014951_59:T>C | C02 | 2526335 | -0.02 | 0 |
| FIRST FLOWERING | 2015 | 4112381 | C02 | 2720764 | 1.710146 | 5.94E-05 |
| LAST FLOWERING | 2015 | 4112381 | C02 | 2720764 | 1.371163 | 1.65E-05 |
| NDVI (14July) | 2015 | 3134538 | C02 | 3021170 | -0.03 | 0 |
| NDVI (27July) | 2015 | 3084367_42:A>T | C03 | 22640779 | 0.01 | 0 |
| NDVI (14July) | 2015 | 3084367_42:A>T | C03 | 22640779 | 0.01 | 0 |
| NDVI (11Aug) | 2015 | 3084367_42:A>T | C03 | 22640779 | 0.01 | 0 |
| BMFresh | 2014 | 7507901 | C03 | 23251374 | 10.20058 | 0.000245 |
| NDVI (14July) | 2015 | 5810998 | C03 | 23251374 | 0.01 | 0 |
| NDVI (27June) | 2014 | 5810998 | C03 | 23251374 | 0.02 | 0 |
| NDVI (20June) | 2014 | 5810998 | C03 | 23251374 | 0.02 | 0 |
| NDVI (27July) | 2015 | 5816633_59:G>A | C03 | 23287237 | 0.01 | 0.01 |

| LOD score | Genotypic variance explained (%) | Multi-trait QTL (Distance from neighboring marker, bp) |
|---|---|---|
| 5.596707 | 18.7124 | |
| 5.960226 | 19.8461 | |
| 3.688185 | 12.38062 | 0 |
| 1.9 | 2.1 | |
| 2.08 | 6.6 | 882670 |
| 2.5 | 3.5 | |
| 1.615175 | 4.877863 | 107119 |
| 2.288815 | 7.36023 | |
| 5.1 | 7.4 | |
| 2.326165 | 7.497341 | 102354 |
| 2.161674 | 6.892731 | |
| 3.92 | 13.19 | |
| 4.226001 | 14.22935 | 194429 |
| 4.783699 | 16.0933 | |
| 6.48 | 21.43 | 300406 |
| 2.66 | 8.71 | |
| 3.63 | 12.17 | |
| 2.57 | 8.4 | |
| 3.610947 | 12.11109 | 610595 |
| 2.37 | 7.65 | |
| 2.55 | 8.33 | |
| 2.56 | 8.34 | |
| 1.86 | 5.79 | |
